# Vaccine-elicitation of cross-group neutralizing protective antibodies to influenza A viruses

**DOI:** 10.1101/2022.02.11.480115

**Authors:** Syed M. Moin, Jeffrey C. Boyington, Seyhan Boyoglu-Barnum, Rebecca A. Gillespie, Gabriele Cerutti, Crystal Sao-Fong Cheung, Alberto Cagigi, John R. Gallagher, Joshua Brand, Madhu Prabhakaran, Yaroslav Tsybovsky, Tyler Stephens, Brian E. Fisher, Adrian Creanga, Sila Ataca, Reda Rawi, Kizzmekia S. Corbett, Michelle C. Crank, Jason Gorman, Adrian B. McDermott, Audray K. Harris, Tongqing Zhou, Peter D. Kwong, Lawrence Shapiro, John R. Mascola, Barney S. Graham, Masaru Kanekiyo

## Abstract

Current influenza vaccines predominantly induce immunity to the hypervariable viral hemagglutinin (HA) head, requiring frequent vaccine reformulation. Conversely, antigenic sites on the conserved HA stem are subdominant and harbor a supersite which is targeted by broadly neutralizing antibodies (bnAbs), making it a prime target for universal vaccines. Here, we show that co-immunization of two stem immunogens derived from influenza A group 1 and 2 HAs elicits cross-group protective immunity and neutralizing antibody responses in mice, ferrets, and nonhuman primates (NHPs). Immunized mice were protected from multiple group 1 and 2 viruses, and all animal models showed broad serum neutralizing activity. A bnAb isolated from an immunized NHP broadly neutralized and conferred protection from viruses including H5N1 and H7N9. Genetic and structural analyses revealed a remarkable convergent evolution between macaque and human bnAbs, illustrating the biophysical constraints for acquiring immunoglobulins with cross-group specificity. Co-immunization of stem immunogens elicits not only group-specific protective immunity, but also cross-group bnAb responses, and represents a step towards broadly protective influenza vaccines.

## MAIN TEXT

Influenza virus causes a substantial global health and economic burden despite the availability of commercial vaccines. Moreover, it poses a unique biological challenge by evading host immune responses through its genetic plasticity leading to antigenic variation, making the virus a constantly moving target. Current influenza vaccines provide narrow-spectrum protection against antigenically closely matched strains, resulting in inconsistent 10–60% vaccine efficacy against symptomatic disease in humans^1^. This also necessitates annual update of the vaccine strains in order to match the predicted circulating viruses. Furthermore, these vaccines are not expected to confer meaningful protection against zoonotic virus strains from animal reservoirs that cause sporadic outbreaks in humans and have pandemic potential.

Current influenza vaccines target the most abundant influenza viral glycoprotein, hemagglutinin (HA). HA is a homotrimeric class I fusion protein which densely decorates virions along with another glycoprotein neuraminidase (NA). To date, 18 phylogenetically different HA subtypes and 11 NA subtypes have been identified in influenza A viruses. These 18 HA subtypes are categorized in 2 distinct groups (group 1 and 2). The HA glycoprotein is expressed as a precursor HA0 which is subsequently cleaved by cellular proteases into HA1 and HA2 (ref. ^2^). This cleavage is required for virus infectivity and HA1 and HA2 are covalently linked by disulfide bonds^3^. HA has two functionally and structurally distinct regions, head and stem, that mediate host cell attachment and viral membrane fusion processes, respectively. The hypervariable head region contains antigenic sites that are highly neutralization sensitive and immunodominant, whereas the stem region is considerably more conserved within each group and recognized by broadly neutralizing antibodies (bnAbs) that are typically less potent than head-directed monoclonal antibodies (mAbs). These antibodies also utilize Fc-mediated effector functions such as antibody dependent cell-mediated cytotoxicity to broaden protection against diverse influenza strains^4–6^. Multiple broadly neutralizing monoclonal antibodies have been isolated from humans that are capable of neutralizing different group 1 or group 2 subtypes or viruses from both groups^7,8^. Hence, one major strategy for universal influenza vaccine efforts has been to focus on the HA stem domain to achieve broad protection against seasonal and pandemic influenza viruses without yearly reformulation. We and others have shown previously that rationally-designed immunogens based on the HA stem elicit broadly cross-reactive antibody responses and confer heterosubtypic protection against H5N1 infection in animal models^9–13^. However, cross-group protection against heterosubtypic group 2 viruses has not been reported for HA stem-based immunogens. Recently, parallel approaches have been pursued to generate group 2 HA stem immunogens based on H3 and H7 (refs. ^14,15^). Although these stem immunogens elicited intra-group cross-reactive antibody responses and homotypic protective immunity in small animals, the induction of bnAb responses that resemble human bnAbs has not been demonstrated. The group 2 H3 and H7 HA stem displayed on ferritin nanoparticles were shown to activate B cells expressing B cell receptors of unmutated common ancestors (UCAs) for human cross-group bnAbs representing two multi-donor classes, suggesting that these immunogens have potential to activate such precursor cells in humans. Joyce et al.^7^ showed that H5 HA immunization in human clinical trials resulted in bnAbs against group 1 and group 2 influenza viruses. Co-crystal structures with HA demonstrated recognition of overlapping epitopes on the HA stem by these human classes of bnAbs using germline genes and convergent sequence motifs. One such antibody class utilized V_H_6-1 and D_H_3-3 (V_H_6-1+D_H_3-3) recognizing the HA stem and avoiding the conserved HA1 glycans of the respective group 1 and group 2 subtypes. In each case, a somatically mutated CDR H3 residue was inserted into the conserved Trp21_HA2_ pocket in the hydrophobic groove of HA2. Small animal models like mice and ferrets have been widely used for studying vaccine responses and antibody mediated protection. However, these animal models do not recapitulate the human immune repertoire due to the lack of sequence homology with human immunoglobulin genes. Non-human primates (NHPs) are evolutionarily close to humans and can generate immune responses similar to humans, which makes them a preferred model for vaccine studies especially when focused on specific antibody lineages. Moreover, NHPs have recently been shown to elicit bnAbs against influenza viruses^16^. Although NHPs lack V_H_ genes possessing the hydrophobic CDR H2 tip as seen in the human immunoglobulin germline V_H_1-69, the V_H_ gene most commonly used by group 1 HA stem-recognizing human bnAbs^17^, there are some with human resemblance, including those similar to genes contributing to the V_H_6-1+D_H_3-3 class of human multi-donor bnAbs that can facilitate HA stem recognition in monkeys^16^.

Here, we extend our ability to design additional group 2 stabilized HA stem trimers displayed on ferritin nanoparticles using the HA sequence of an avian H10N8 virus (H10ssF) which causes sporadic infections in humans. We found that the resulting H10ssF elicited broadly protective antibody responses against challenge with multiple heterosubtypic group 2 viruses in mice. We also explored co-immunization with the HA stem of group 1 (H1ssF) and group 2 (H10ssF) to elicit cross-group protective immunity in mice, ferrets, and NHPs. Further, we isolated and characterized vaccine-elicited monoclonal antibodies for their specificity and protective efficacy. We found that a combined immunization of group 1 and group 2 HAssFs conferred not only contiguous group-specific neutralizing and protective antibody responses but also cross-group reactive bnAb responses that recognized the stem supersite in a manner reminiscent of the human multi-donor V_H_6-1+D_H_3-3 class bnAbs, providing a step towards developing broadly protective influenza vaccines.

### Design and characterization of H10 HA stabilized stem nanoparticle (H10ssF)

We have previously developed a method to structurally stabilize group 2 HA stem such as H3 and H7 in native-like trimers and display on self-assembling ferritin nanoparticles (H3ssF and H7ssF, respectively) as potential immunogens for eliciting antibody responses to the highly conserved HA stem^14^. While H3ssF and H7ssF induced stem-directed protective immune responses to homotypic viruses in mice, neither of them had shown to elicit heterosubtypic protective immunity that spans across multiple group 2 subtype viruses. To overcome this hurdle of conferring heterosubtypic protective immunity, we sought to improve the group 2 stem immunogen by redesigning it with different HA subtype sequences. To identify a candidate group 2 HA that has a better cross-reactive potential, we determined the conservation of primary sequences as well as solvent-exposed residues among different group 2 HA stems (Supplementary Fig. 1a,b). We found that the H10 stem is uniquely positioned between H3 and H7 phylogenetically and possesses slightly higher sequence and solvent-exposed residue identities both to H3 and H7 than those between H3 and H7 (Supplementary Fig. 1c,d). We hypothesized that the H10-based group 2 stem immunogen might have a better antigenic surface to elicit broadly cross-reactive antibody responses. We employed the same design principle to stabilize the H10 HA stem and generate H10ssF using A/Jiangxi-Donghu/346/2013 (H10N8) as previously described^14^. The self-assembled H10ssF was readily purified from the culture supernatant of transfected mammalian cells which gave a distinct peak around 1.2 MDa corresponding to a size of fully assembled particles on size exclusion chromatography (Supplementary Fig. 2a), with monomeric subunits corresponding to ~45 kDa as predicted (Supplementary Fig. 2b). The purified H10ssF showed homogeneous and well-formed particles when analyzed by negative stain electron microscopy (nsEM) (Supplementary Fig. 2c). A reference-free 2D class averaging of nsEM dataset resulted in particles with spherical cores of ~12 nm diameter and regularly spaced spikes of ~7–8 nm in height, consistent with the design and previously reported group 2 HAssFs^14^. Single-particle cryo-electron microscopy (cryo-EM) reconstruction of the full H10ssF particle resulted in a 4.8 Å resolution structure that revealed structural elements of the ferritin core and the spikes of HA-stem trimers (Fig. 1a, Supplementary Fig. 2d-g). Although the ferritin core could be further resolved through focused refinement, the density for the stem trimers became increasingly diffuse, suggesting their flexibility relative to the core granted by the short 3-amino acid (SGG) linker. Low-pass filtering the density map to 8 Å was able to resolve the stem domain and its angular orientation, while also suggesting the magnitude of the stem domain flexibility. Dynamic light scattering (DLS) showed homogeneous particles with a radius of ~10 nm (Supplementary Fig. 3a). We measured the thermostability of H10ssF by differential scanning fluorometry (DSF) and found that H10ssF was equivalent or more thermally stable than other group 2 HAssFs or group 1 H1ssF^9,14^ (Supplementary Fig. 3b). We next evaluated the antigenic integrity of H10ssF by assessing binding of cross-group, stem-specific, conformation-dependent bnAbs FI6v3 (ref. ^18^), MEDI8552 (ref. ^19^), CT149 (ref. ^20^), 315-53-1F12 (ref. ^8^), group 2 stem-specific CR8020 (ref. ^21^) and 315-24-1E07 (ref. ^8^), and group 1 stem-specific CR6261 (ref. ^22^) and 315-02-1H01 (ref. ^8^) by ELISA. Analogous to H3ssF and H7ssF, H10ssF was recognized by all the stem-directed mAbs except for the group 1-specific mAbs, CR6261 and 315-02-1H01 (Fig. 1b, Supplementary Fig. 3c). We also measured the binding kinetics of the antigen-binding fragment of antibody (Fab) of MEDI8852, CT149 and CR8020 to H10ssF by biolayer interferometry (BLI) and found that H10ssF bound all three bnAbs with high affinities (*K*_D_) of 0.4–4 nM, that were equivalent or of higher affinity than the corresponding H10 HA trimer or other group 2 HAssF (Supplementary Fig. 3d). To evaluate the ability of H10ssF to activate B cells expressing B-cell receptors (BCRs) of the inferred UCA of human bnAbs, we measured Ca^2+^ flux by flow cytometry using recombinant Ramos B cells as described previously^23,24^. We found that H10ssF but not H1ssF activated the recombinant Ramos B cells expressing BCR of 16.a.26 UCA which engages group 2 HA stem (multi-donor V_H_1-18 Q-x-x-V class)^7,14^ (Supplementary Fig. 4a). In contrast, B cells expressing a group 1-specific BCR were activated by H1ssF but not by H10ssF (Supplementary Fig. 4b). These results demonstrate that H10ssF displays conformationally and antigenically intact trimeric headless HA stem spikes that are capable of engaging and activating B cells expressing BCR of human bnAb precursors.

**Figure 1.**
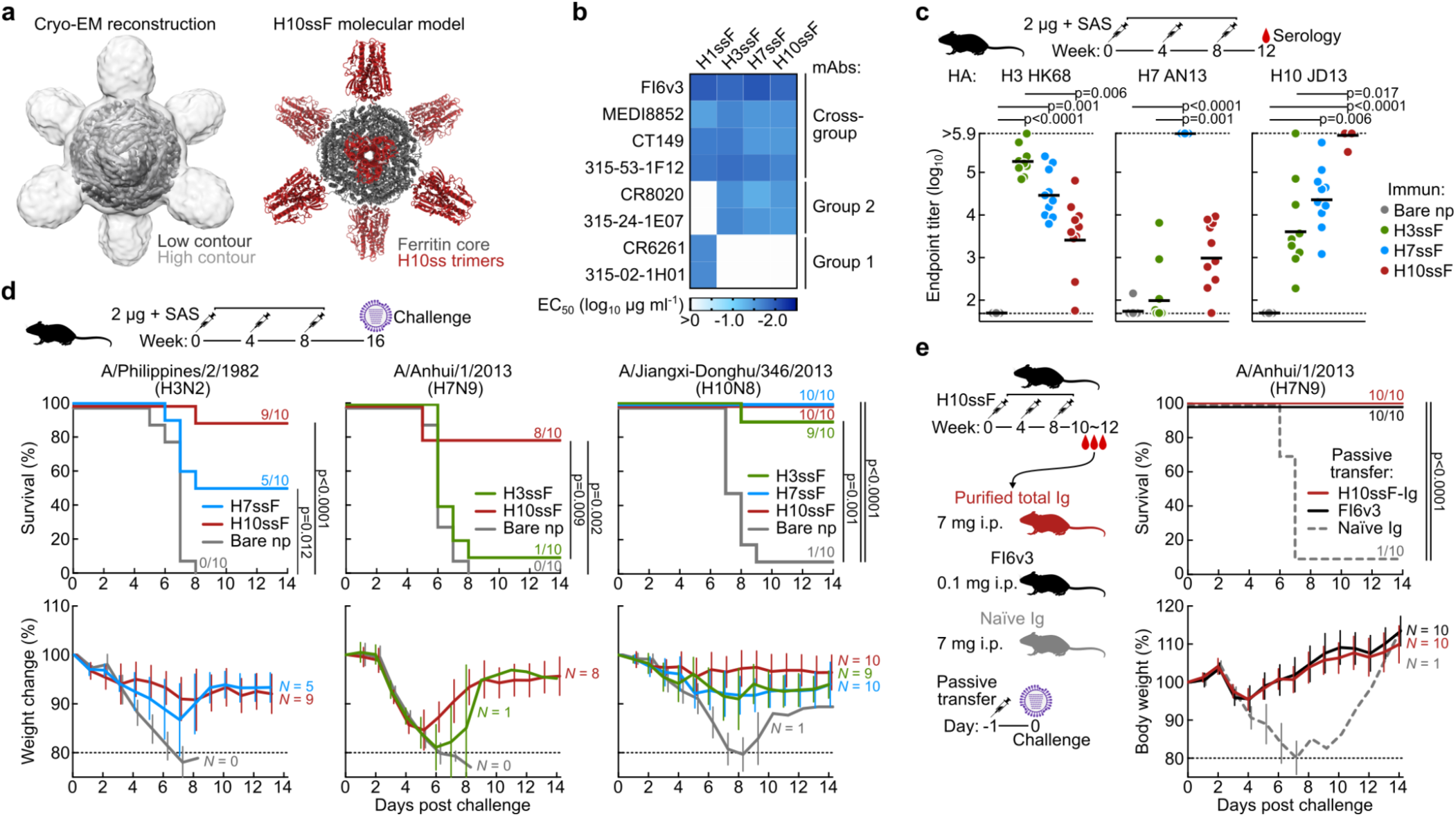
Stabilization, expression and characterization of Group 2 HA-stem ferritin nanoparticles, their immunogenicity and protection efficacy in mice. **a,** Single-particle cryo-EM structure of H10ssF. 3D reconstruction density map of H10ssF (left). The outer contour in light gray was low-pass filtered to 8 Å to reveal the HA-stem trimers, while the inner contour in dark gray indicates the 4.8 Å map of the full complex, contoured to show the ferritin core.. Molecular model of the designed H10ssF showing different structural domains (right). Ferritin core and H10ss spikes are colored in gray and dark red, respectively. **b,** Antigenicity of HAssF determined by ELISA of HA stem-directed mAbs. mAbs FI6v3, MEDI8852, CT149, and 315-53-1F12 recognize both group 1 and group 2 HAs. mAbs CR8020 and 315-24-1E07 recognize group 2 HAs; whereas, mAbs CR6261 and 315-02-1H01 recognize group 1 HAs. **c,** Immunogenicity of group 2 HAssF in BALB/c mice. Mice were immunized 3 times with 2 μg H3ssF, H7ssF, or H10ssF and sera was analyzed at week 12. ELISA antibody titers to A/Hong Kong/1/1968 HA (H3 HK68), A/Anhui/1/2013 HA (H7 AN13), and A/Jiangxi-Donghu/346/13 HA (H10 JD13) are shown. Each dot represents an individual animal and the horizontal bars indicate the geometric mean of each immunization group (N = 10). Horizontal dotted lines denote higher and lower limits of detection. Statistical analysis was performed by Kruskal-Wallis test with Dunn’s multiple comparison post-hoc test. **d,** Protective efficacy of group 2 HAssF in mice. BALB/c mice immunized with H3ssF, H7ssF, or H10ssF (N = 10) were experimentally infected intranasally with either A/Philippines/2/1982 (H3N2), A/Anhui/1/2013 (H7N9), or A/Jiangxi-Donghu/346/2013 (H10N8) viruses 8 weeks post the last immunization. **e,** Passive transfer experiment of purified hyperimmune Igs followed by experimental H7N9 infection. Igs (7 mg) of H10ssF-immunized or naïve mice were passively given to naïve BALB/c mice (N = 10) intraperitoneally 24 h prior to infecting with A/Anhui/1/2013 (H7N9). mAb FI6v3 (5 mg kg^-1^) was used as positive control. Multiple comparisons of Kaplan-Meier curves were performed by the log-rank test with Bonferroni correction (**d,e**).

### Group 2 H10ssF elicits broad heterosubtypic protective immunity in mice

To assess the immunogenicity of H10ssF and compare it with other group 2 HAssF, we immunized mice with H3ssF, H7ssF, or H10ssF in the presence of adjuvant (Sigma Adjuvant System (SAS)). All three immunogens were immunogenic and elicited antibody responses that were cross-reactive to more than one group 2 HA subtype although the highest antibody titers were observed against HAs matched to the immunogen subtype (Fig. 1c). When we measured virus neutralizing activity using the HA/NA pseudotyped lentiviral reporter assay^25^, as expected, we observed serum neutralizing activity against the pseudovirus with matched HA/NA subtype (Supplementary Fig. 5a). There was low but detectable neutralization activity against heterosubtypic pseudoviruses in H7ssF- and H10ssF-immunized mice (Supplementary Fig. 5a). To evaluate protective efficacy of the group 2 HAssF, immunized mice were challenged with a lethal dose of A/Philippines/2/1982 (H3N2), A/Anhui/1/2013 (H7N9) or A/Jiangxi-Donghu/346/2013 (H10N8) influenza viruses. Nearly all of the control mice (97% or 29 of 30 mice) receiving bare ferritin nanoparticles succumbed to infection and were euthanized; in contrast, mice immunized with H10ssF exhibited substantial protection against all 3 viruses tested (93% or 27 of 30 mice survived overall) (Fig. 1d). While both H3ssF and H7ssF were previously demonstrated to confer protection against homologous subtype viruses^14^, they also provided a near-complete (90–100%) protection to heterosubtypic H10N8 virus (Fig. 1d). In contrast, H3ssF and H7ssF conferred only minimal to partial protection against heterosubtypic H7N9 and H3N2 viruses, respectively (10% and 50%, respectively) (Fig. 1d). These results illustrate that H10ssF induces a broader protective immunity against group 2 subtype viruses than either H3ssF or H7ssF in mice. To better understand the basis for this stem-directed protective immunity to group 2 viruses, we tested the specificity of both binding and neutralizing antibodies by using an HA variant which possesses an added N-linked glycan at the epicenter of the canonical stem supersite (Δstem HA) so that it blocks access of vast majority of stem-directed bnAbs^26^. Binding antibody titers of sera from H10ssF immunized mice to Δstem H10 HA were significantly lower than to the wild-type H10 HA (p = 0.0028). Antibody titers to heterosubtypic H7 HA were almost completely abrogated when the Δstem mutation was introduced (Supplementary Fig. 5b), suggesting that H10ssF immunization elicited antibodies that target the canonical stem supersite surrounding the Trp21_HA2_ hydrophobic groove^27^ and those antibodies were responsible for heterosubtypic cross-reactivity. Similarly, sera from H7ssF-immunized mice showed reduced binding to both homologous and heterologous HAs in the presence of the Δstem mutation (Supplementary Fig. 5b). Importantly, the serum neutralizing activity of H10ssF-immunized mice was not substantially diminished by the excess Δstem HA protein unlike the wild-type HA which fully depleted the activity (Supplementary Fig. 5c). In addition, serum immunoglobulins (Igs) purified from H10ssF-immunized mice (H10ssF-Igs) reacted with H10 and H7 HA, and the binding was significantly lower to Δstem H10 HA and nearly abolished to Δstem H7 HA as seen in immune sera (Supplementary Fig. 5d). The H10ssF-Igs neutralized homologous H10N8 or heterosubtypic H3N2 and H7N9 pseudoviruses (Supplementary Fig. 5d). To confirm that the heterosubtypic group 2 protection is antibody-mediated, H10ssF-Igs were passively administered to naïve mice prior to infecting them with a lethal dose of H7N9 virus. Mice receiving the H10ssF-Igs were fully protected from the virus with no measurable weight loss (Fig. 1e), while all mice receiving Igs purified from naïve mice succumbed to infection. This result shows that the heterosubtypic group 2 protection can be achieved by stem-directed antibodies elicited by H10ssF.

### Co-immunization of group 1 and group 2 HAssF confers cross-group heterosubtypic immunity

To achieve the goal of eliciting protective immune responses to influenza A viruses carrying either group 1 and group 2 subtype HAs, we evaluated a cocktail vaccine approach consisting of H1ssF and H10ssF. Groups of mice, ferrets and cynomolgus macaques were immunized with a cocktail of H1ssF and H10ssF (termed herein H1+10ssF) with a squalene oil-in-water emulsion adjuvant (AddaVax®). Immunization of H1+10ssF elicited HA-binding and neutralizing antibody responses to several group 1 and group 2 subtype viruses in mice (Fig. 2a,b), ferrets (Fig. 2c) and nonhuman primates (NHPs) (Fig. 2d). The levels of antibody responses to group 1 viruses were comparable to that induced by H1ssF alone (Fig. 2a–c), and likewise, the responses to group 2 viruses were comparable to that induced by H10ssF alone (Fig. 2a–c). These findings confirm that the co-immunization of group 1 and group 2 HAssF does not cause antigenic competition between the two HAssF immunogens. Importantly, co-immunization also conferred substantial protection in immunized mice from heterosubtypic H5N1 and H3N2 lethal virus challenges (Supplementary Fig. 6a,b). Although heterosubtypic virus neutralizing antibody responses in mice and ferrets could be measured by pseudotyped lentiviral neutralization assays, serum neutralizing activity in immunized NHPs were also readily detectable in influenza reporter virus-based microneutralization assays (Fig. 2d)^28^. Mouse and ferret immune sera neutralized all 5 pseudotyped viruses that represent group 1 and group 2 human influenza viruses, whereas NHP immune sera did not show detectable neutralization against the H3N2 virus (Fig. 2b–d). These results suggest that immunizations with H1+10ssF can elicit broadly neutralizing serum antibody responses that are capable of neutralizing multiple group 1 and group 2 subtype viruses although the breadth may be more limited in NHPs.

**Figure 2.**
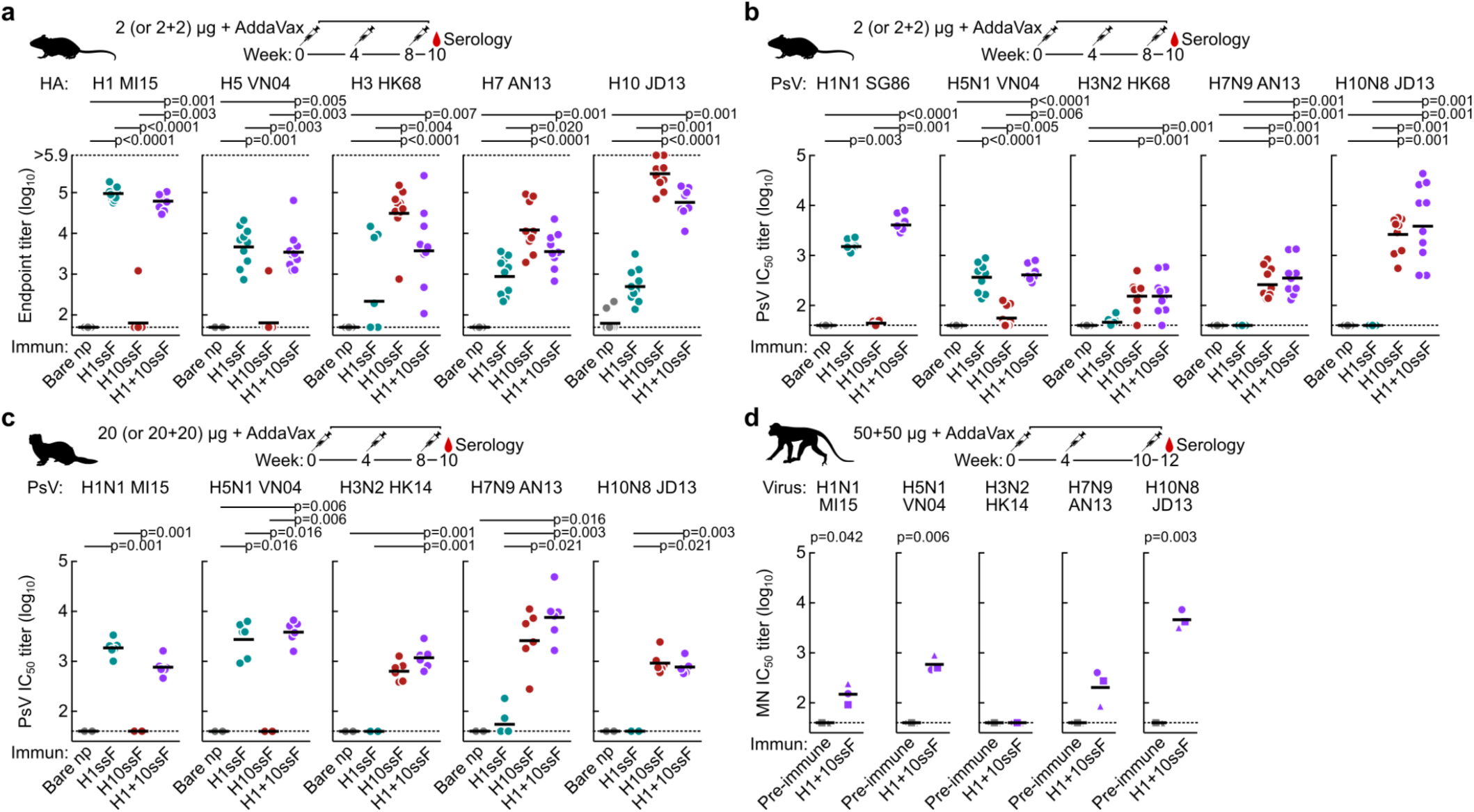
Co-immunization of group 1 and group 2 HAssF induced broadly cross-reactive antibody responses. **a,** Cross-reactive HA-binding antibody responses to multiple group 1 and group 2 HAs. Sera (N = 10) were collected after 3 immunizations with H1ssF, H10ssF, H1+10ssF, or bare nanoparticles. ELISA antibody responses were assessed for A/Michigan/45/2015 (H1 MI15), A/Vietnam/1203/2004 (H5 VN04), H3 HK68, H7 AN13, and H10 JD13 HA. **b,c,** Neutralizing antibody responses to pseudotyped lentiviruses expressing multiple group 1 and group 2 HAs in mice (**b**) and ferrets (N = 6) (**c**). Pseudotype neutralization assays were performed by using HA and NA from A/Singapore/1/1986 (H1N1 SG86), H1N1 MI15, H5N1 VN04, H3N2 HK68, H7N9 AN13, and H10N8 JD13. **d,** Neutralizing antibody responses in NHPs. Cynomolgus macaques (N = 3) were immunized with H1+10ssF. Serum neutralizing activity as measured by the reporter-based microneutralization assay^28^. H1N1 MI15, H5N1 VN04, A/Hong Kong/4801/2014 (H3N2 HK14), H7N9 AN13, and H10N8 JD13 reporter influenza viruses were used. Different symbols are used to identify individual animals. Black horizontal lines indicate geometric mean titers for each respective group. Horizontal dotted lines denote higher and lower limits of detection (**a**) or lower limit of detection (**b–d**). Statistical analysis was performed by Kruskal-Wallis test with Dunn’s multiple comparison post-hoc test (**a**–**c**) or by paired *t*-test (**d**).

### Induction of heterosubtypic and cross-group HA-specific B cells in nonhuman primates

To better understand the basis for cross-reactive antibody responses observed in NHPs immunized with H1+10ssF, we assessed the B cell responses to multiple HAs in peripheral blood mononuclear cells (PBMCs) by flow cytometry. Two weeks after the second immunization (week 6), B cells binding to homologous H1 MI15 HA and H10 JD13 HA were readily detectable (Fig. 3a, Supplementary Fig. 7). Frequencies of IgG^+^ B cells recognizing homologous H1 and H10 HA were substantially increased after the third immunization (Fig. 3b). At week 12, a large fraction (~50%) of the H1^+^ IgG^+^ B cells showed cross-reactivity to H5 VN04 HA and likewise, approximately half of the H10^+^ B cells bound H7 AN13 HA; whereas, cross-reactivity between H10 and H3 HK14 HAs was markedly lower than H7 (Fig. 3c). B cells that recognize both group 1 (H1) and group 2 (H10) HAs were rare yet detectable (Fig. 3a). We next assessed the kinetics for the development of cross-reactive B cell responses in the animal (M08145, circle) which had the highest cross-group HA-reactive B cells. The frequency of B cells recognizing homologous HAs (i.e., H1 and H10) were low but detectable as early as one week after the first H1+10ssF immunization (week 1) (Fig. 3d). Frequencies of both homologous H1^+^ and H10^+^ B cells were remarkably increased after the second and the third immunizations (week 5 and week 11, respectively). A disproportional increase of H5^+^ B cells compared to H1^+^ only B cells was observed after the second and the third immunization, indicating that selective induction of cross-reactive B cells occurred after each H1+10ssF booster (Fig. 3d). Notably, B cells cross-reacting with both H1 and H10 (cross-group) emerged after the second immunization, while the relative frequency of these cells were substantially enriched after the third immunization (Fig. 3e). These results demonstrate that a cocktail of H1ssF and H10ssF is capable of inducing a diverse repertoire of cross-reactive HA-specific B cells including some that cross-react with both group 1 and group 2 HAs.

**Figure 3.**
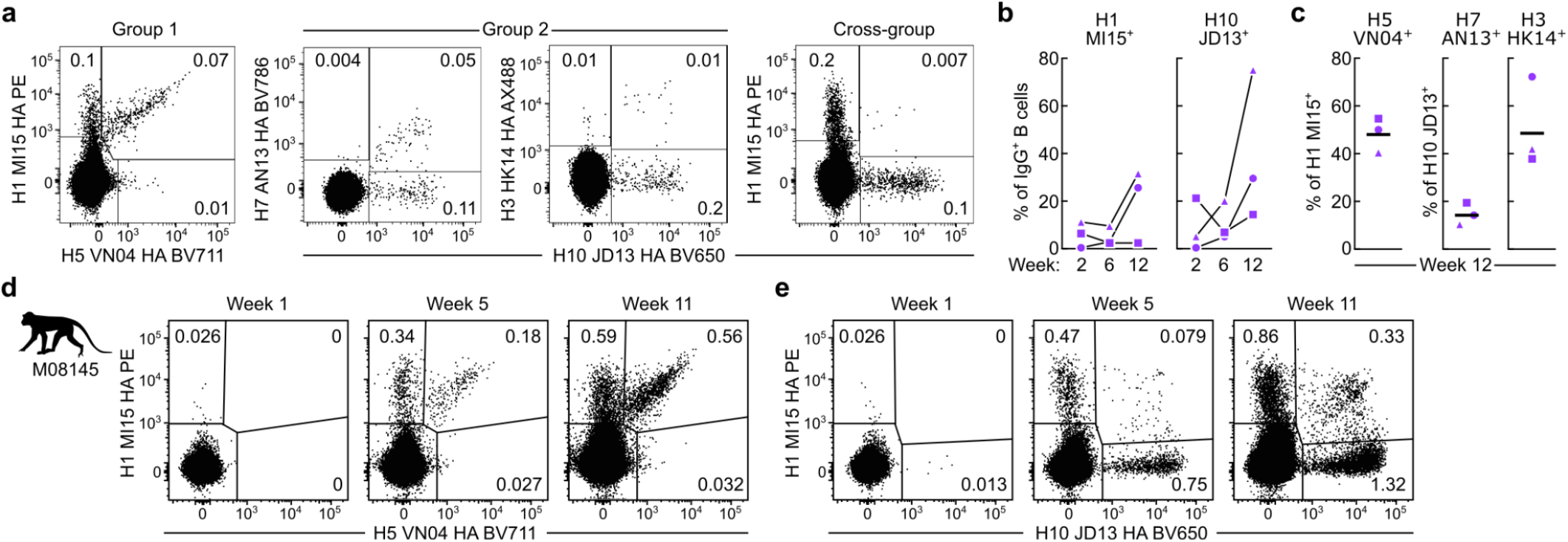
Induction of cross-reactive HA-specific B cells in NHPs. **a,** Detection of HA-specific B cells by flow cytometry. PBMCs from NHPs (N = 3) at two weeks after the second immunization were analyzed by using fluorescently-labeled HA trimer probes. **b,** Frequencies of homologous H1^+^ and H10^+^ B cells in PBMCs. Percentage of H1^+^ or H10^+^ among IgG^+^ B cells are shown. **c,** Frequencies of cross-reactive B cells. Percentage of H5 positivity among H1+ B cells (left), and percentages of H7^+^ or H3^+^ among H10^+^ B cells (right) are shown. Horizontal lines indicate the geometric mean of a group. **d,** Kinetics of cross-reactive B cell responses in macaque M08145. PBMCs taken at one week after each immunization were analyzed by flow cytometry with H1 MI15, H5 VN04, and H10 JD13 HA trimer probes.

We next investigated the functionality of cross-group HA-specific B cells by performing B cell sorting and sequencing. We single-cell sorted H1^+^ H10^+^ B cells from PBMCs (week 12) of the animal M08145 and sequenced their immunoglobulin (Ig) genes. Recombinant antibodies were produced by co-expressing Ig heavy and light chain genes encoding Ig variable domains recovered from the single-cell sorted B cells. Among the mAbs derived from cross-group HA-specific B cells, we found an antibody clone 789-203-3C12 that bound multiple subtype HAs and neutralized a wide range of influenza viruses in the reporter-based microneutralization assays (Fig. 4a,b). Binding affinities of the 789-203-3C12 Fab to various subtypes of HA varied ranging from 6.6 to 534 nM (Fig. 4a). Although the mAb 789-203-3C12 did not show detectable binding to H3 HA nor neutralized H3N2 viruses, it bound and neutralized multiple group 1 and group 2 HA subtype viruses (Fig. 4b). 789-203-3C12 was highly protective against both group 1 (A/California/04/2009 (H1N1) and H5N1 VN04) and group 2 (H7N9 AN13 and H10N8 JD13) virus infections when the antibody was given prophylactically to naïve mice (Fig. 4c). These results establish that the co-immunization of H1+10ssF can induce robust B cell responses that are cross-reactive to heterosubtypic HAs not only within either group 1 or group 2, but also across the two HA groups. In addition, the mAb 789-203-3C12 isolated from H1+10ssF-immunized NHP showed cross-group neutralizing activity as well as protective efficacy to multiple group 1 and group 2 subtype influenza viruses.

**Figure 4.**
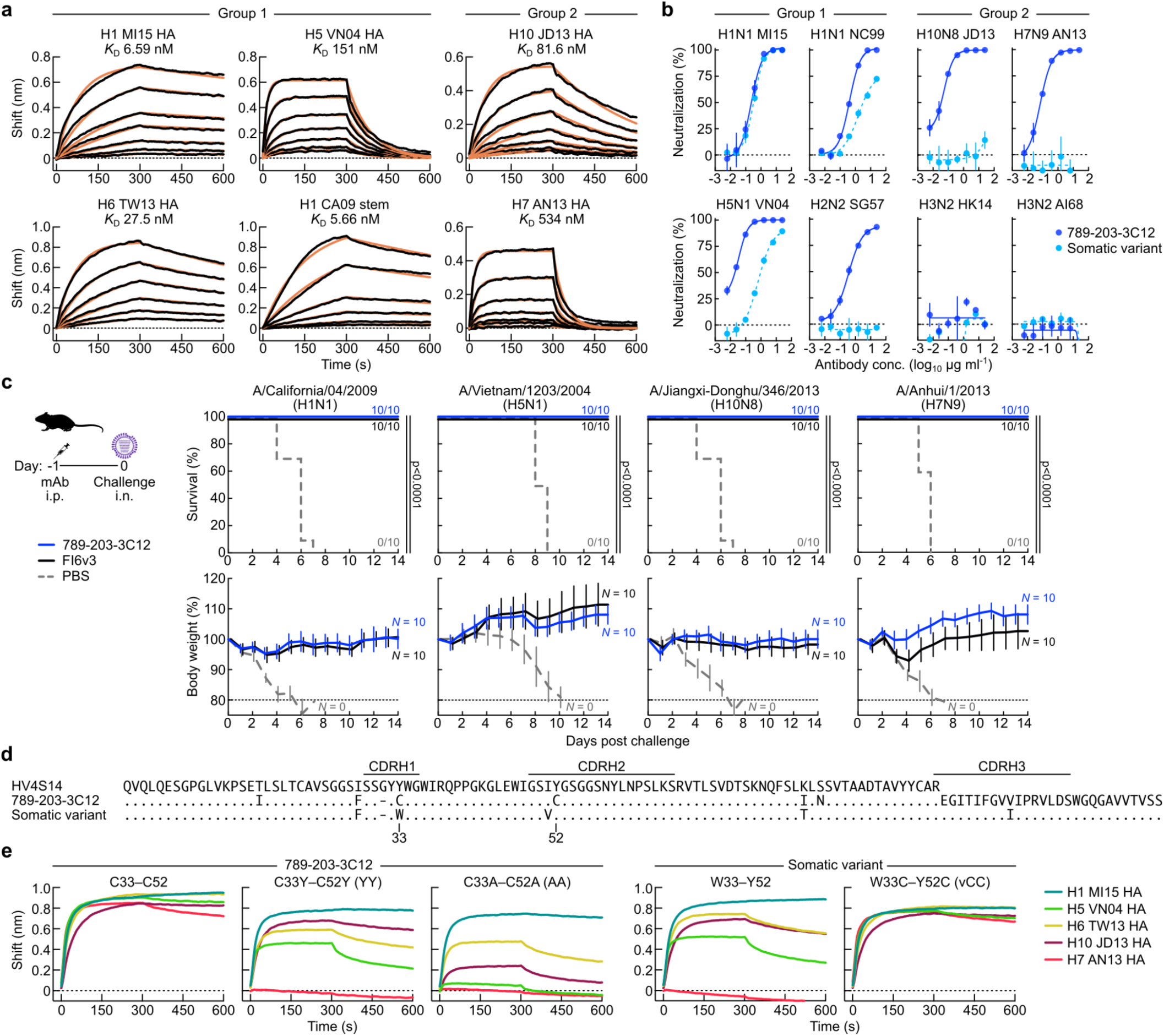
Neutralizing breadth, potency, and protective efficacy of NHP mAb 789-203-3C12. **a,** HA-binding kinetics of Fab 789-203-3C12 determined by BLI. Recombinant HA of multiple group 1 and group 2 subtypes or the stabilized HA stem were immobilized on biosensors. Experimental datasets were fitted with the binding equations describing a 1:1 interaction to obtain apparent affinities. **b,** Neutralization profile of mAb 789-203-3C12 and its somatic variant antibody. Neutralization curves were generated by the reporter-based microneutralization assay^28^. **c,** Protective efficacy of mAb 789-203-3C12 in mice. 789-203-3C12 or control FI6v3 antibodies were passively administered (10 mg kg^-1^) to recipient BALB/c mice (N = 10) intraperitoneally 24 h prior to infecting with multiple group 1 and group 2 subtype viruses. Multiple comparisons of Kaplan-Meier curves were performed by the log-rank test with Bonferroni correction. **d,** Sequence alignment of the germline-encoded HV4S14 heavy chain, 789-203-3C12, and its somatic variant. Dots denote identical residues to the HV4S14 whereas the residues that are different from the HV4S14 are indicated. CDR loop regions are shown by horizontal lines (Kabat). **e,** Binding profile of mAb 789-203-3C12 and the somatic variant antibody with their CDR H1 and CDR H2 variants. HA-binding was measured using BLI. Recombinant HAs were immobilized on biosensors. All the mAbs used in the assay were tested as IgG.

### Mutational and immunogenetic characterization of the vaccine-elicited cross-group bnAb 789-203-3C12

To study the molecular basis for broad cross-group neutralization, we characterized immunogenetic composition of 789-203-3C12. The V_H_ region of 789-203-3C12 is highly similar to the *M. fascicularis IGHV4S14* gene (96.2%), suggesting it acquired a pair of Cys substitutions within CDR H1 and H2 regions (Tyr33Cys and Tyr52Cys, respectively) alongside a few other substitutions in its framework regions during somatic hypermutation (SHM) (Fig. 4d). Among the sequences of single-cell sorted B cells, we found a somatic variant heavy chain sequence clonally related to 789-203-3C12 which did not possess the double Cys substitutions in its CDR H1 and H2 loops (Fig. 4d). Interestingly, when the recombinant antibody was produced using the somatic variant heavy chain paired with 789-203-3C12 light chain, the resultant mAb had comparable neutralizing activity against H1N1 MI15 virus (Fig. 4b); however, this somatic variant mAb had a much reduced neutralizing activity against heterologous pre-pandemic H1N1 (A/New Caledonia/20/1999) and heterosubtypic H5N1 VN04 viruses. Moreover, its neutralizing activity against the group 1 H2N2 or the group 2 subtype viruses was completely abolished (Fig. 4b). To test whether the Cys substitutions were critical for its binding and neutralization breadth and potency, we generated two variant 789-203-3C12 heavy chains by substituting the Cys residues to either the germline-encoded Tyr (Cys33Tyr/Cys52Tyr, YY) or the smaller side-chain Ala (Cys33Ala/Cys52Ala, AA). Both the YY and AA mutant mAbs produced by pairing with the 789-203-3C12 light chain bound H1 HA in a manner analogous to the parental 789-203-3C12 (Cys33/Cys52, CC) but showed reduced or no binding to H5, H6, H7 and H10 HAs (Fig. 4e). Interestingly, introducing the double Cys substitutions in the CDR H1 and H2 of the somatic variant heavy chain (Trp33Cys/Tyr52Cys, vCC) was sufficient to convert the binding profile of the resultant mAb (vCC) from that of the somatic variant to resemble 789-203-3C12 (Fig. 4e). These results suggest that the acquisition of double Cys in the CDR H1 and H2 loops during SHM was critical to the optimal cross-group breadth and neutralization potency of the mAb 789-203-3C12.

### Structure of the cross-group bnAb 789-203-3C12 in complex with HA

To understand the structural basis for HA recognition by 789-203-3C12, we determined the structure of 789-203-3C12 Fab in complex with recombinant stabilized HA trimer^29^ of H1N1 virus (A/Solomon Islands/3/2006, SI06) to 3.85 Å resolution by single-particle cryo-EM (Fig. 5a, Supplementary Fig. 8). While 60,632 particles showed C3 symmetry, the dataset for the HA–Fab complex structure was collected based on the 116,041 particles which showed C1 symmetry (Supplementary Table 1). 789-203-3C12 bound to the canonical HA stem supersite, with HA1 and HA2 subunits providing 15% (153 Å^2^) and 85% (837 Å^2^), respectively, of a total buried surface area (BSA) of 990 Å^2^ (Supplementary Table 2). The heavy chain dominated the binding interface and contributed 75% of the 1,066.32 Å^2^ paratope surface (Supplementary Table 2). In the CDR H3, Arg100f formed a salt bridge with the HA residue Asp19_HA2_, and Val100c interacted with the Val18_HA2_ through hydrophobic contacts (Fig. 5b, Supplementary Table 3). Phe100 at the tip of the CDR H3 bound in a hydrophobic groove formed by residues Trp21_HA2_, Ile45_HA2_, and Ile48_HA2_ (Fig. 5b). Overall, CDR H3 contributed ~51% of the total paratope surface (Supplementary Tables 4,5). As expected, Cys33 and Cys52 formed an intra-chain disulfide bond, with well-resolved electron density (Supplementary Fig. 8e), linking CDR H1 and CDR H2 (Fig. 5b). This CDR H1-H2 disulfide bond provided perfect shape complementarity to accommodate the loop consisting of the HA2 residues 16-21, and helped orient the Val18_HA2_ and Asp19_HA2_ to interact with CDR H3 (Fig. 5b). In addition, CDR H2 residues Gly55 and Lys64 formed hydrogen bonds and salt bridges with Tyr34_HA2_ and Asp146_HA2_, respectively (Fig. 5b, Supplementary Table 4,5). The light chain residues Arg50, Ser91, and Thr92 formed hydrogen bonds with Gln42_HA2_, Asn46_HA2_, and Thr49_HA2_ (Fig. 5b, Supplementary Tables 6,7).

**Figure 5.**
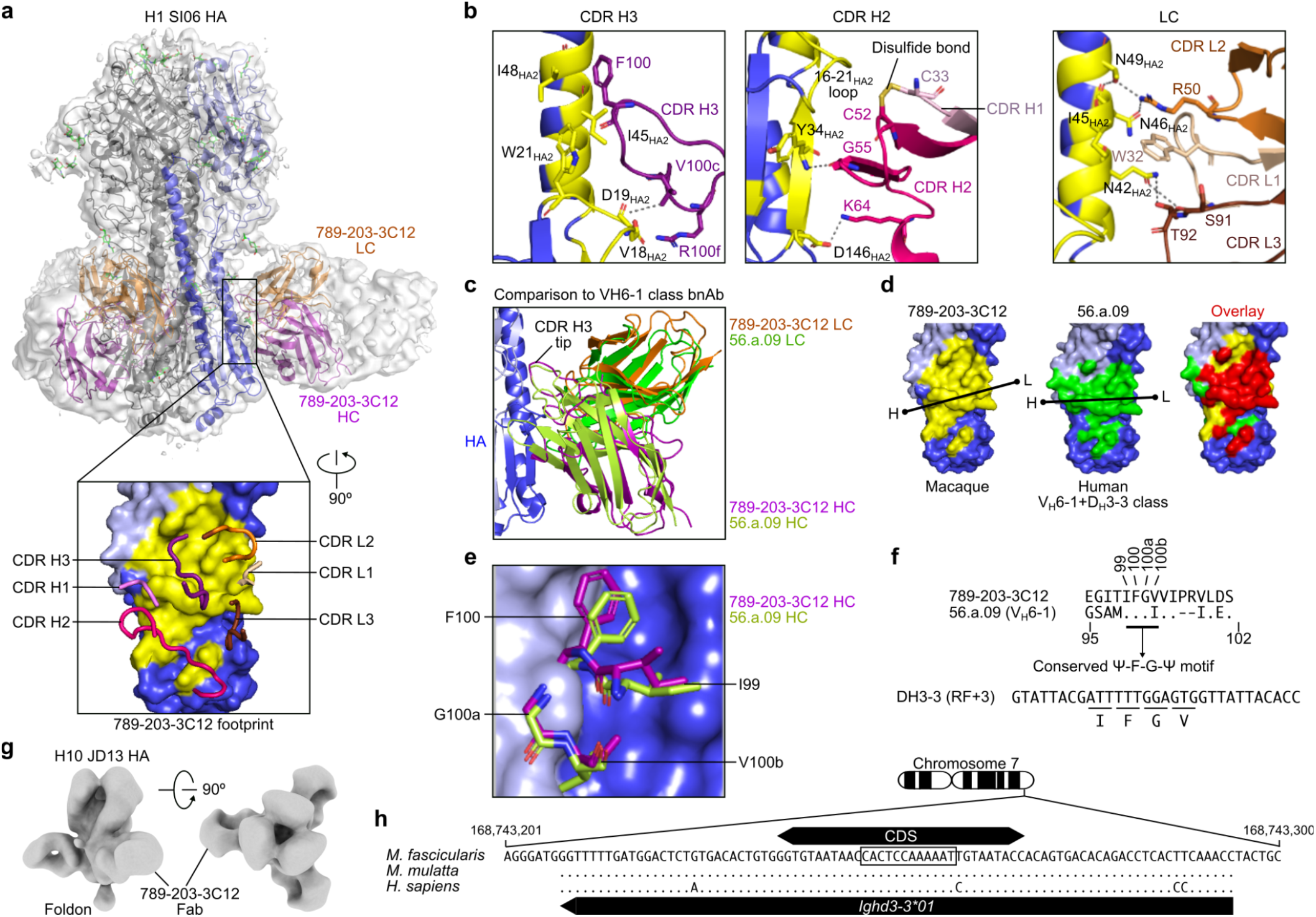
Structural characterization of NHP mAb 789-203-3C12. **a,** Overall structure of the 789-203-3C12 Fab bound to the H1 SI06 HA trimer. The HA trimer is shown in gray, with the protomer with bound antibody is colored blue. HA1, HA2, and antibody heavy and light chains are depicted as ribbons colored light blue, blue, purple, and orange, respectively. A semitransparent surface representation of the entire complex is also show in gray. A 90° rotated view of the interface with interacting CDR loops shown as worms and HA as surface representation (inset). The epitope of 789-203-3C12 is colored in yellow. **b,** Detailed interactions of 789-203-3C12 CDR loops with HA. The formation of a disulfide bond between CDR H1 and H2 loops are shown in yellow (middle panel). Hydrogen bonds and salt bridges are indicated as dashed lines. **c,** Comparison of the binding mode of 789-203-3C12 antibody with the human V_H_6-1+D_H_3-3 class bnAb human 56.a.09 on HA. **d,** The footprints for 789-203-3C12 (yellow) and 56.a.09 (green) are mapped onto the HA surface. A black line indicates the orientation of the antibody defined by the line connecting Cα atoms of heavy chain Cys22 (dot with letter H) and light chain Cys23 (dot with letter L). The overlapping epitope was colored in red (right). **e,** Structural arrangement of the “Ψ-F-G-Ψ” motif in the 789-203-3C12 and 56.a.09. **f,** Sequence alignment of the CDR H3 of 789-203-3C12 and 56.a.09. Human D_H_3-3 gene is shown (bottom). **g,** Negative stain EM 3D reconstruction of 789-203-3C12 Fab bound to the H10 JD13 HA trimer. **h,** Sequence alignment of the (putative) D_H_3-3 genes of cynomolgus macaque (*M. fascicularis*), rhesus macaque (*M. mulatta*), and human (*H. sapiens*). Ideogram of *M. fascicularis* is shown on top. CDS based on human D_H_3-3 is shown as a double arrow box. The nucleotides encoding the IFGV are boxed. Orientation of the human *Ighd3-3*01* gene is indicated (bottom).

We noted that the macaque-derived 789-203-3C12 recognized the HA stem similar to the convergent V_H_6-1+D_H_3-3 class of human bnAbs^7^. One such human V_H_6-1+D_H_3-3 class antibody, 56.a.09 (PDB 5K9K), shared a very similar approach to the HA stem, where the heavy and light chains were superimposable with nearly identical orientations (Fig. 5c). The epitopes for 789-203-3C12 and 56.a.09 showed a high degree of overlap, sharing 21 residues (Fig. 5d). Although the structures of CDR H1, H2 and CDR L1, L2, and L3 are substantially different between 789-203-3C12 and 56.a.09, the CDR H3 of both antibodies contained the conserved Ile-Phe-Gly-Val/Ile (Ψ-F-G-Ψ) motif, enabling both antibodies to interact with HA in a similar fashion (Fig. 5e,f). Additionally, nsEM analyses of the 789-203-3C12 Fab in complex with H10 HA (A/Jiangxi-Donghu/346/2013) confirmed that the Fab bound the group 2 HA stem in a manner analogous to human V_H_6-1+D_H_3-3 class bnAbs (Fig. 5g). Structures of group 2 HA–Fab complex (MEDI8852–H7 HA (PDB 5JW3) and 56.a.09–H3 HA (PDB 5K9K) fit snugly into the reconstructed nsEM density of 789-203-3C12–H10 complex (Supplementary Fig. 9), further highlighting the similarity between macaque and human crossgroup bnAbs. While the conserved Ile-Phe-Gly-Val/Ile motif in the V_H_6-1+D_H_3-3 class bnAbs is entirely encoded by the human IGHD3-3 gene, this same motif was completely conserved in the CDR H3 of 789-203-3C12 (Fig. 5f). Due to the lack of complete immunoglobulin loci of cynomolgus macaque genome in public databases, we were unable to correctly assign the *Ighd* gene used by 789-203-3C12. However, we found that the nucleotide sequence corresponding to the D segment within the *M. fascicularis* (cynomolgus macaque) genome (GCA_011100615.1) was identical to the D_H_3-3 gene of *M. mulatta* (rhesus macaque) (GenBank MF989451.1). It is also highly homologous to the human D_H_3-3 gene, and encodes the core “Ile-Phe-Gly-Val” motif (Fig. 5h). These facts suggest that the corresponding D_H_ gene (putative *Ighd3-3*) is also present in the macaque genomes and is utilized by influenza HA stem-directed bnAbs.

## DISCUSSION

The ongoing global SARS-CoV-2 pandemic underscores the difficult public health challenges encountered when fighting against highly contagious respiratory pathogens. Vaccines remain the most effective countermeasure to slow down transmission as demonstrated by the highly efficacious vaccines against SARS-CoV-2. Even though the vaccines are partially protective, if they were deployed early or prior to the pandemic, they could have a considerable positive impact on the epidemic curve^30^. This is particularly relevant to influenza as it is not possible to accurately predict the next pandemic virus with the available technologies and research tools. Therefore, the development of vaccines for influenza pandemic preparedness that are effective against unknown pandemic viruses and their deployment immediately after a new virus emerges would be immensely valuable for maintaining global public health. Headless HA stem immunogens could also be combined with conventional split vaccines or with full-length HA immunogens to improve the breadth of immunity and could reduce the need for yearly vaccination. Alternatively, a vaccine that induces broad immunity against group 1 and 2 influenza A viruses, could be stockpiled and available for deployment during the early stages of a pandemic before strain-matched vaccines can be made available. Our study demonstrated the feasibility of such a vaccine approach using a cocktail of the group 1 and group 2 headless HA stem-nanoparticles^9,14^ and showed that it could elicit broadly neutralizing and protective immune responses against a wide range of influenza A viruses in three different animal models. Markedly, this cocktail vaccine approach induced not only within-group (group 1 or group 2) cross-reactive antibodies but also the cross-group bnAbs in NHPs. To the best of our knowledge, this is the first study reporting the vaccine-elicitation and isolation of cross-group bnAbs resembling a public human bnAb clonotype in NHPs, providing a key step to the development of vaccines that elicit bnAbs in humans. To date, such cross-group bnAbs induced by infections and/or vaccinations in humans are rarely found^7,8,18,19,31,32^.

For more than a century, small animal models have been indispensable to gain insights on fundamental biological processes and study immunological mechanisms. These animal models have been widely used in preclinical studies for evaluating vaccine immunogenicity; however, results obtained from these models do not always recapitulate complex immune responses in humans. For example, in humans, the predominant bnAb responses to group 1 HA stem are largely constrained by the usage of V_H_1-69 genes which do not exist in nonhuman animal species^33,34^. To better address this issue, transgenic animals that carry human immune receptor genes have been developed and utilized to reproduce human-like immune responses^35–38^. Despite the advances in the development of a variety of humanized transgenic and transchromosomic mice over the last few decades, it is still exceedingly difficult to model the complexity of human immune responses. NHPs and in particular macaques are the preferred animal species for vaccine evaluation due to their evolutionary proximity to humans. While NHPs do not possess the human V_H_1-69 gene which is by far the most prevalent gene utilized for group 1 HA stem-directed bnAbs, they use a different set of V_H_ genes (i.e., V_H_3S5 and V_H_3S25) to target the same stem epitope^16^. Analogous to humans, these group 1 HA stem-directed antibodies were highly dominant and genetically constrained public clonotypes in NHPs. In addition to the group 1 HA stem-directed antibodies, NHPs have been shown to generate group 2 HA stem-directed antibodies following H3 stem-nanoparticle immunizations^16^. Noteworthily, the NHP antibodies to the group 2 HA stem target the membrane proximal epitope similar to the human group 2 HA stem-directed antibodies, such as CR8020 and CR8043 (refs. ^21,39^). Despite the fact that NHPs were capable of generating human-like bnAb responses to either group 1 or group 2 HA stem following infection or vaccination^16,40–42^, there are no bnAbs that broadly neutralize both group 1 and group 2 influenza viruses reported to date. In the present study, we identified the NHP bnAb 789-203-3C12 as the first example of such cross-group bnAbs.

Although the bnAb 789-203-3C12 was isolated from a cynomolgus macaque, the antibody was surprisingly similar to the human V_H_6-1+D_H_3-3 class of bnAbs^7,19^. The canonical CDR H3 motif of this class of bnAbs is entirely encoded by the D_H_3-3 gene in humans. We found the putative homolog of human D_H_3-3 gene in both rhesus and cynomolgus macaques’ genomes that were nearly identical to the human D_H_3-3. While it is likely that this D_H_ gene has been conserved in those macaque species for completely irrelevant reasons, it is astounding to observe the convergent solution to recognize the influenza HA stem between humans and macaques by utilizing this gene out of many D_H_ genes with a particular reading frame. It is unclear at the moment whether or not the V_H_ gene (V_H_4S14) poses an immunogenetic bottleneck in macaques like the human V_H_6-1 gene does for the V_H_6-1+D_H_3-3 public clonotype. We also have limited understanding on why bnAb 789-203-3C12 does not neutralize H3N2 viruses and if this limitation is common amongst NHP cross-group bnAbs. Further antibody discovery efforts would be needed to address these questions and leverage the NHP model for studying the germline gene-endowed immunological processes in response to the HA stem.

Although we and others have shown that the HA stem-directed broadly cross-reactive antibody responses can be elicited in multiple animal models through active immunization of designed immunogens^9,14,15,43–45^, the potential impacts of preexisting influenza immunity on vaccine-elicitation of the stem-directed antibody responses have not been rigorously studied. A recent natural history study uncovered a substantial impact of early life influenza exposures on protective immunity against heterosubtypic influenza virus infections in humans^46^. The observed protective immunity appeared to be HA group-specific and determined by the subtype of influenza virus that the individual was first exposed to in their early childhood (H1N1, H2N2, or H3N2). This “immunological imprint” introduces birth year-dependent bias in immune repertoire and may influence group preference of the vaccine-induced immunity, posing another layer of difficulty for achieving universal influenza immunity in humans^47^. Whether a designed vaccine can override and reprogram the imprinted group-biased immunity in humans remains to be determined, although there are vaccine candidates in advanced clinical trials that show some promise in phase 1 human clinical trials by eliciting the HA stem-directed antibody responses in adults with preexisting influenza immunity^48–50^. In addition, the building block of our stem-nanoparticles is encoded by a single gene and its assembly is precise and efficient, making it a candidate for genetic vaccine delivery approaches such as mRNA and recombinant viral vectors. These vaccine modalities have been pivotal for combating the ongoing SARS-CoV-2 pandemic and are anticipated to play major roles in other vaccines onward. These modalities along with innovative new adjuvants would offer not only manufacturing options and dose sparing but also different immunological properties including durability and cellular immunity. The present study provides a proof-of-concept for the vaccine-elicitation of broad cross-group protective immunity and would foster future studies addressing outstanding challenges such as preexisting immunity and durability. Furthermore, both group 1 HA stem- and group 2 HA stem-nanoparticles reported herein have entered phase 1 clinical trials (NCT03814720 and NCT04579250, respectively) to evaluate safety and immunogenicity in humans. Results from these trials would inform the feasibility of the HA stem-based vaccine strategies for eliciting bnAb responses and reshaping the immune repertoire in humans in the face of preexisting influenza immunity.

## METHODS

### Structure based immunogen design

The complete sequence for A/Jiangxi-Donghu/346/2013 (H10N8) including the signal sequence, was obtained from the GISAID accession number EPI497477. The crystal structure of A/Jiangxi-Donghu/346/2013 (H10N8) HA (PDB ID 4QY0) was used as a template for modeling using the “C” version of HAssF as previously described^14^. Loops were modeled with the program LOOPY^51^, superpositions were performed using UCSF Chimera^52^ or LSQMAN^53^, energy minimization was performed using RELAX^54^ and the free energy changes from point mutations were calculated using the program DDG_MONOMER^55^ from the ROSETTA suite. PyMOL (Schrodinger, LLC) and UCSF Chimera^52^ were used for detailed visualization of structures and structural figure generation.

### Expression and purification of immunogens, antibodies and probes

The H3ssF, H7ssF and H10ssF sequences each incorporated the “C” version of group 2 HAssF as described previously^14^. The HAssF nanoparticles, H3ssF, H7ssF, H10ssF and H1ssF, monoclonal antibodies (mAbs) FI6v3 (ref. ^18^), CT149^20^, CR8020 (ref. ^21^), CR6261 (ref. ^22^), MEDI8852 (ref. ^19^), 315-53-1F12, 315-54-1G07, 315-04-1D02, 315-09-1B12 (ref. ^8^), 789-203-3C12 (this study), and recombinant foldon-trimerized HA ectodomain trimers of H1 MI15 (A/Michigan/45/2015), H5 VN04 (A/Vietnam/1203/2004), H3 HK14 (A/Hong Kong/4803/2014), H7 AN13 (A/Anhui/1/2013), H10 JD13 (A/Jiangxi-Donghu/346/2013), and H1 SI06 (A/Solomon Islands/3/2006) were produced by transient transfection of the corresponding plasmid(s) in Expi293 cells (Life Technologies) using Expifectamine293 transfection kit (ThermoFisher Scientific) according to manufacturer’s instruction. All HAssF and mAbs were purified with *Galanthus nivalis* lectin (GNA-gel, EY Laboratories) and protein A (rmp Protein A Sepharose, Cytiva), respectively as previously described^9^. Recombinant HA proteins were purified by Ni-NTA through C-terminal hexa-histidine tag. HAssF and HA proteins were further purified by size exclusion chromatography using Superose 6 increase 10/300 GL and Superdex 200 increase 10/300 GL columns, respectively (Cytiva). To produce Fabs, MEDI8852, CR8020, and 789-203-3C12 IgG was digested with endoproteinase Lys-C (New England Biolabs) overnight at room temperature. The Fab of CT149 was produced by digesting CT149 IgG with an engineered HRV-3C cleavage site^56^ using HRV-3C enzyme (MilliporeSigma) overnight at RT. Upon completion, the reactions were quenched by adding protease inhibitor cocktail (MilliporeSigma), and Fc and Fabs were separated by passing through protein A column.

### Animals and immunizations

All animal experiments were reviewed and approved by the Institutional Animal Care and Use Committee of the Vaccine Research Center, NIAID, NIH. All animals were housed and cared for in compliance with the pertinent US National Institutes of Health regulations and policies and American Association for Accreditation of Laboratory Animal Care. Female BALB/cJ mice (N = 10) aged between 6 and 8 weeks (Jackson Laboratory) were immunized with 2 μg of H3ssF, H7ssF, H10ssF or H1ssF formulated with the Sigma adjuvant system at weeks 0, 4, and 8 with 100 μl via intramuscular route, given as 50 μl into each hind leg. Two weeks post second (week 6) and final immunization (week 10), sera were collected for immunological assays. For experimental virus challenge studies, mice were inoculated intranasally with a 10 × 50% lethal dose (LD_50_) of H3N2 A/Philippines/2/1982, 20 × LD_50_ of H7N9 A/Anhui/1/2013, 10 × LD_50_ of H10N8 A/Jiangxi-Donghu/346/13 or 25 × LD_50_ of H5N1 A/Vietnam/1203/2004 virus at BioQual. Infected animals were monitored daily for 14 days post-challenge for their weight change and symptoms. For ferrets (female and male ~6 months old fitch ferrets, TripleF Farm), animals (N = 6) were given 20 μg of HAssF formulated with SAS in 500 μl via intramuscular route at weeks 0, 4 and 8. Serum samples were collected 2 weeks after each immunization for serological assays. For NHP study, cynomolgus macaques (*M. fascicularis*, N = 3) were immunized with a mixture of 50 μg each of H1ssF and H10ssF formulated with AddaVax (Invivogen) in 1 ml via intramuscular route at weeks 0, 4, and 10. Blood samples were periodically collected after each immunization for immunological assays. For immunoglobulin (Ig) passive transfer studies, serum Igs were purified from H10ssF-immunized mice (combined 2-4 weeks after the third immunization) by using protein A/G (Pierce Thermo Scientific). Seven mg of purified heperimmune Igs or control Igs or 100 μg (5 mg kg^-1^) of FI6v3 IgG were given via intraperitoneal route (i.p.) to naïve recipient BALB/cAnNHsd mice (N = 10) aged between 4 and 6 weeks (Envigo) 24 h prior to the experimental intranasal challenge with 20 × LD_50_ of H7N9 A/Anhui/1/2013 virus. Infected animals were monitored daily for 14 days post-challenge for their weight change and symptoms. For mAb passive transfer studies, 200 μg of mAb 789-203-3C12 or FI6v3 were given i.p. to naïve recipient mice as described above. Twenty-four hours later, mice were experimentally challenged with 20 × LD_50_ of H1N1 A/California/04/2009, 25 × LD_50_ of H5N1 A/Vietnam/1203/2004, 20 × LD_50_ of H7N9 A/Anhui/1/2013, or 10 × LD_50_ of H10N8 A/Jiangxi-Donghu/346/13 virus at BioQual. Infected animals were monitored daily for 14 days post-challenge for their weight change and symptoms.

### ELISA binding assays

ELISA was performed to assess anti-HA antibodies in response to HAssF immunization. The plates were coated with 200 ng well^-1^ of recombinant full-length HA-foldon proteins and incubated overnight at 4°C. The plates were then blocked with 5% skim milk in PBS at 37°C for 1 h. The plates were incubated with 4-fold serially diluted mAbs with dilution series starting at 10 ng ml^-1^ or immune sera dilutions starting at 1:50. Appropriate horseradish peroxidase (HRP)-conjugated secondary antibodies (Southern Biotech) were added and incubated for 1 hr followed by a colorimetric detection assay utilizing 3,3′,5′,5-tetramethylbenzidine (TMB) substrate (KPL). The colorimetric reaction was stopped by addition of 1 M H_2_SO_4_ and the absorbance recorded at 450 nm (OD_450_). Serum endpoint titers were recorded as serum dilution resulting in 4-fold increase in OD_450_ value above the background level.

### Biolayer interferometry binding assays

The biolayer interferometry binding assays were done as previously described^14,57^. All biosensors were hydrated in PBS prior to use. CR9114 (ref. ^31^) was used at 10 μg ml^-1^ in PBS containing 1% bovine serum albumin (PBS-BSA) to coat AHC biosensors (fortéBio) to capture H10ssF. The CR9114-coated biosensors were dipped in H10ssF (10 μg ml^-1^ in PBS-BSA). After equilibrated for 60 s in PBS-BSA, the biosensors were then dipped in a 2-fold dilution series of MEDI8852, CT149, or CR8020 Fab for 300 s, followed by incubation in PBS-BSA for 300 s to allow for dissociation. For 789-203-3C12, recombinant HAs (10 μg ml^-1^) were immobilized on HIS1K biosensors through hexa-histidine tag. After briefly equilibration in PBS-BSA, the biosensors were dipped in a 2-fold dilution series of 789-203-3C12 Fab for 300 s followed by dissociation in PBS-BSA for 300 s. All assay steps were performed at 30°C with agitation set at 1,000 rpm in the Octet HTX instrument (fortéBio). Correction to subtract non-specific baseline drift was carried out by subtracting the measurements recorded for a bare sensor. Data analysis and curve fitting were carried out using Octet analysis software (v9). Experimental data were fitted with the binding equations describing a 1:1 interaction. Global analyses of the complete data sets assuming binding was reversible (full dissociation) were carried out using nonlinear least-squares fitting allowing a single set of binding parameters to be obtained simultaneously for all concentrations used in each experiment.

### Negative stain electron microscopy analysis

Purified H3ssF, H7ssF, and H10ssF were diluted to 0.02 mg ml^-1^ with buffer containing 10 mM HEPES, pH 7.0, and 150 mM NaCl and adsorbed onto glow-discharged carbon-coated copper mesh grids for 15 s. The grids were washed with the buffer and stained with 0.7% uranyl formate. Images were obtained at a magnification of 50,000 semi-automatically on an FEI Tecnai T20 transmission electron microscope (TEM) operated at 200 kV, equipped with a 2,048-by 2,048-pixel Eagle charge-coupled-device (CCD) camera using SerialEM^58^. The pixel size was 0.44 nm and particles were picked automatically using software developed in-house. Reference-free 2D classification was performed using SPIDER^59^ and Relion 1.4 (ref. ^60^). For 3D reconstruction of the H10 JD13 HA trimer in complex with 789-203-3C12 Fab, the purified components were mixed at a molar ratio of 1.1 Fab to 1 HA protomer, briefly incubated on ice, and negatively stained as described above. A dataset containing 144 micrographs was collected using the same TEM at a nominal magnification of 100,00x, corresponding to a pixel size of 0.22 nm, and 4746 particle images were selected using e2boxer from EMAN2 (ref. ^61^) and extracted into 128-by 128-pixel boxes. After 2D classification in Relion 3.0, selected 2D class averages were used to generate an ab-initio 3D model in EMAN2. This was followed by 3D classification and 3D autorefinement in Relion with imposed C3 symmetry. The final dataset contained 1173 particles, and the resolution at the Fourier shell correlation threshold of 0.5 was 24.8 Å.

### Cryo-EM structure of H10ssF

Purified H10ssF was applied to glow discharged Quantifoil R 2/1 300 mesh grids (Ted Pella, Redding) at a concentration of 1 mg ml^-1^. Using a Leica EM GP plunge freezer (Leica Microsystems), the grid was blotted for 1 s at 37°C and 68% humidity, then plunge-frozen in liquid ethane. Plunged grids were loaded into a Polara electron microscope (ThermoFisher Scientific) operated at 300 keV, equipped with a Falcon 2 direct electron detector (ThermoFisher Scientific), and EPU data acquisition software (ThermoFisher Scientific). Total of 4,027 micrographs, dose fractionated into 7 frames, were collected at a magnification of 78,000 ×, with 1.43 Å^2^ pixel^-1^ and a total dose of 47 e^-^ pixel^-1^. Movie frames were aligned using MotionCor2 (ref. ^62^), and the contrast transfer function for each micrograph was fitted using CTFFIND4 (ref. ^63^). Total of 262 particles were manually picked to generate class averages with RELION2 (ref. ^64^), which were used for autopicking of 693,018 particles by Gautomatch v0.56 (https://www2.mrc-lmb.cam.ac.uk/download/gautomatch-056/). Further curation and alignment of particle sets was performed with RELION2. Particles were 2D classified in batches of 30,000-60,000, culling overlapping particles and false-positive particle picks from the analysis. For 3D analysis, an initial model of a solid sphere of diameter 220 Å was used, combined with an outer spherical mask of diameter 325 Å. Iterative 3D refinement of 118,473 particles against the model using “gold-standard” refinement of 2 independent half-maps as implemented in RELION2 generated a map at a resolution of 4.8 Å, using a 0.143 FSC cutoff and post-processed with an ad-hoc B-factor of −300. While antigenic spikes on H10ssF were observed during initial low-resolution refinement, their density became unresolved as the refined resolution improved. The location of the antigenic spikes could again be visualized by low-pass filtering the final map back to 8 Å resolution.

### Cryo-EM structure of Fab-HA complex

The sample for cryo-EM analysis of 789-203-3C12 Fab in complex with stabilized HA trimer of H1N1 virus (A/Solomon Islands/3/2006, SI06) was produced by mixing the Fab and HA trimer in a 6:1 molar ratio, with a final trimer concentration of 0.2 mg ml^-1^, followed by incubation on ice for 1 hr. The protein complex was purified by a Superdex 200 16/60 column in a buffer containing 5 mM HEPES pH 7.5 (Life Technologies) and 150 mM NaCl (Quality Biological, Inc.). To prevent aggregation during vitrification, 0.005% (w/v) n-dodecyl β-D-maltoside was added to the sample prior to plunge freezing. Cryo-EM grids were prepared by applying 2 ml of sample to a freshly glow-discharged carbon-coated copper grid (CF 1.2/1.3 300 mesh); the sample was vitrified in liquid ethane using a Vitrobot Mark IV with a wait time of 30 s and a blot time of 3 s. Cryo-EM data were collected using the Leginon software ^65^ installed on a Titan Krios electron microscope operating at 300 kV, equipped with a Gatan K3-BioQuantum direct detection device. The total dose was fractionated for 3 s over 60 raw frames. Motion correction, CTF estimation, particle extraction, 2D classification, ab initio model generation, 3D refinements and local resolution estimation for all datasets were carried out in cryoSPARC 2.15 (ref. ^66^). The final 3D reconstruction was obtained using non-uniform refinement with C1 symmetry. HA trimer density was modeled using PDB entry 6FYT^67^, as initial template. The heavy chain variable region of 789-203-3C12 Fab was modeled using PDB entry 4FQQ^68^ while the light chain variable region was modeled using PDB entry 5OD8 (ref. ^69^). Automated and manual model building were iteratively performed using real space refinement in Phenix^70^ and Coot^71^, respectively. Half maps were provided to the Resolve Cryo-EM tool in Phenix to support manual model building. Geometry validation and structure quality assessment were performed using EMRinger^72^ and Molprobity^72,73^. Map-fitting cross correlation (Fit-in-Map tool) and figures preparation were carried out using PyMOL and UCSF Chimera^52^ and ChimeraX^52,74^. A summary of the cryo-EM data collection, reconstruction and refinement statistics is shown in Supplementary Table 1.

### Differential scanning fluorimetry

Thermal unfolding of purified HAssF in PBS was measured by nano differential scanning fluorometry (nDSF) using Prometheus NT.48 instrument (NanoTemper technologies). Thermal unfolding curves are plotted slope (Δ(F350 nm/F330 nm)/ΔT) against temperature (°C) generated from fluorescence measurements at 350 and 330 nm. Thermal transition midpoint temperature (Tm) was calculated from the curves and reported.

### Dynamic light scattering

Dynamic light scattering (DLS) measurements were performed using a DynaPro Plate Reader II (Wyatt Technology). Protein samples of 0.5 mg ml^-1^ were centrifuged at 13,000 rpm in a tabletop refrigerated microfuge at 4°C for 45 minutes prior to aliquoting 30 μl of each into clear flat bottom 384-well plate (Corning). Measurements for each sample were taken at 25°C in the laser auto-attenuation mode and a wavelength of 830 nm from a total of 10 data acquisitions. DLS data was processed using Dynamics 7.8.2 software using a Rayleigh spheres model.

### Influenza pseudovirus and microneutralization assays

Pseudoviruses used for the assay were based on lentivirus vectors encoding the luciferase gene and expressing influenza HA and NA as previously described^26^. Replication-restricted reporter influenza viruses used in this study were previously described^28^. Animal serum was treated with receptor-destroying enzyme, RDE (Denka Seiken) for 37°C, 16 h followed by RDE denaturation at 56°C for 1 h. For pseudovirus neutralization assay, a four-fold serial dilution of mouse sera starting at 1:40 were mixed with either H3N2 (A/Hong Kong/1/1968, HK68), H7N9 (A/Anhui/1/2013, AN13), H10N8 (A/Jiangxi-Donghu/346/2013, JD13), H1N1 (A/Michigan/45/2015, MI15) or H5N1 (A/Vietnam/1203/2004, VN04) pseudoviruses for 30 min at room temperature (RT). The serum-pseudovirus mixture was added to previously plated 293A cells (10,000 cells well^-1^ in a 96-well plate) in triplicate. After 2 h fresh Dulbecco modified Eagle medium (DMEM) supplemented with 10% fetal bovine serum (FBS), 2 mM glutamine, and 1% penicillin–streptomycin was added. Cells were lysed at 72 h, and luciferase assay (Promega) was performed according to manufacturer’s protocol. Luciferase activity was measured as RLU at 570 nm on a SpectramaxL (Molecular Devices). For influenza neutralization assay, a four-fold serial dilution starting at 1:40 (serum samples) or 25 μg ml^-1^ (mAbs) were mixed with one of these replication-restricted reporter viruses: H1N1 (A/Michigan/45/2015, MI15); H3N2 (A/Hong Kong/4801/2014, HK14); H5N1 (A/Vietnam/1203/2004, VN04); H7N9 (A/Anhui/1/2013, AN13); H10N8 (A/Jiangxi-Donghu/346-2/2013, JD13) for 60 min at RT. The serum-virus mixtures were added to previously plated MDCK-SIAT1 cells stable expressing influenza PB1 gene (3,000 cells well^-1^ in a 384-well plate with transparent bottom) in quadruplicate. Plates were scanned at 18-22 h post infection and fluorescence foci were counted using Celigo Image Cytometer (Nexcelom). In Celigo operation and analysis software v5.0, Target 1 protocol was used to detect and count fluorescent foci^28^. IC_50_ titers were calculated considering uninfected cells as 100% and cells transduced with only virus as 0% neutralization.

### B cell activation assays

The B cell activation assay was performed as described before^14,23^. Briefly, to test the ability of HAssF to engage BCR and induce calcium flux, Ramos cells expressing inferred unmutated common ancestor (UCA) sequence of bnAbs as IgM on their surface were utilized. These cells were stained with fura red (ThermoFisher Scientific) in serum free medium at room-temperature for 30 min. Cells were washed and resuspended in serum free medium, incubated at 37°C for 5 min followed by acquisition on a FACSymphony interfaced to the FACSDiva software (BD Biosciences). Cells were acquired without the nanoparticle (baseline) followed by acquisition in presence of the nanoparticle and acquisition for 180 s. The anti-IgM was used as positive control to assess maximal calcium flux induction measured as change in emission by bound versus unbound fura red to calcium over time. The inferred UCA for 01.a.44 was used as control that only recognizes group 1 HAs and H1ssF served as control for 16.a.26 UCA which only binds to group 2 HAs.

### PBMC isolation, staining and single-cell sorting

Cryopreserved peripheral blood mononuclear cells (PBMCs) were used. A panel of antibodies were used to stain the cells and gate for particular cell subsets. Antibodies to CD16 (clone 3G8), CD3 (clone SP34-2), CD56 (clone B159), IgM (clone G20-127), IgG (clone G18-145) (BD Biosciences); CD20 (clone 2H7), CD14 (clone M5E2) (Biolegend); and CD19 (clone J3-119) (Beckman Coulter) were used for cell surface staining. Various group 1 (H1 MI15, H5 VN04) and group 2 (H3 HK14, H7 AN13, H10 JD13) full-length ectodomain HA probes with the sialic acid-binding site mutation (Y98F) were prepared as previously described^75^. HA probes were conjugated with fluorescently-labeled streptavidin through biotin ligated to the Avi-tag on each probe. The cells were stained with a panel of antibodies to the surface markers and HA-streptavidin probes. The viability dye 7AAD was used for excluding dead cells. The samples were analyzed on a FACS Aria II (BD Biosciences) and single-cell sorted into 96-well plates. Index sorting was applied to identify specificity of individual cells for multiple HA probes. Data analysis and subsequent flowplots were generated following cell sorting using FlowJo (v10). PCR amplification of immunoglobulin heavy and light chain genes and cloning into expression vectors were performed as previously described^76^. Immunoglobulin gene assignment was performed by using IMGT/HighV-QUEST (http://www.imgt.org).

### Statistical analysis

Statistical analysis was performed with Prism (v9, GraphPad). Specific details of statistical analysis are indicated in the figure legends and the Results section. P-values less than or equal to 0.05 were considered significant. Normality tests were conducted on all data to determine the appropriate statistical test. All statistical tests used were two tailed.

### Reporting summary

Further information on research design is available in the Nature Research Reporting Summary linked to this article.

## Data availability

All data are available in the main text or the supplementary materials. Cryo-EM structures have been deposited in the Electron Microscopy Data Bank (EMDB) under accession code EMD-26044 (H10ssF) and EMD-22302 (789-203-3C12 Fab–HA complex); and in the Protein Data Bank (PDB) under accession code 6XSK (789-203-3C12 Fab–HA complex). Any additional data are available upon reasonable request from the corresponding author.

## Acknowledgements

The authors thank R. Woodward, D. Scorpio, J.-P. Todd, E. McCarthy, A. Taylor, H. Bao, C. Chiedi, M. Dillon, L. Gilman, and G. Sarbadorand (VRC) for help with animal studies; K. Foulds, A. Noe, S.-F. Kao, V. Ficca, N. Nji, and D. Flebbe (VRC) for help with nonhuman primate samples; H. Andersen, N. Jones, and G. Patel (Bioqual) for help with influenza challenge studies; V. Nair and E. Fisher (NIAID) for cryo-electron microscopy data collection for H10ssF; and S. Andrews and members of VRC influenza program and Viral Pathogenesis Laboratory for helpful discussion. This work used the computational resources of the NIH HPC Biowulf cluster (http://hpc.nih.gov) and the Office of Cyber Infrastructure and Computational Biology (OCICB) High Performance Computing (HPC) cluster at the NIAID, NIH.

## Author contributions

Conceptualization: S.M.M., J.C.B., B.S.G., M.K.; Formal Analysis: SM.M., S.B.-B., R.A.G., C.S.C., A.Ca., J.R.G., M.S.P., Y.T., T.Z., A.K.H.; Investigation: S.M.M., J.C.B., S.B.-B., R.A.G., G.C., C.S.C., A.Ca., J.R.G., J.B., M.S.P., Y.T., T.S., B.E.F., A.Cr., S.A., R.R., K.S.C., J.G., A.B.M., A.K.H., T.Z., P.D.K., L.S., J.R.M., B.S.G., M.K.; Resources: A.Cr.; Writing—original Draft: S.M.M., M.K.; Writing—review and editing: All authors; Supervision: A.B.M., A.K.H., T.Z., P.D.K., L.S., J.R.M., B.S.G., M.K.; Project Administration: M.C.C.; Funding Acquisition: A.K.H., P.D.K., L.S., J.R.M., B.S.G., M.K.

## Additional Information

**Supplementary information** The online version contains supplementary material available at XXX.

**Correspondence and requests for materials** should be addressed to Masaru Kanekiyo.

## SUPPLEMENTARY INFORMATION

Supplementary Fig. 1–9

Supplementary table 1–7

