## Supplementary materials for "Vaccine-elicitation of cross-group neutralizing protective antibodies to influenza A viruses"

Moin et al.

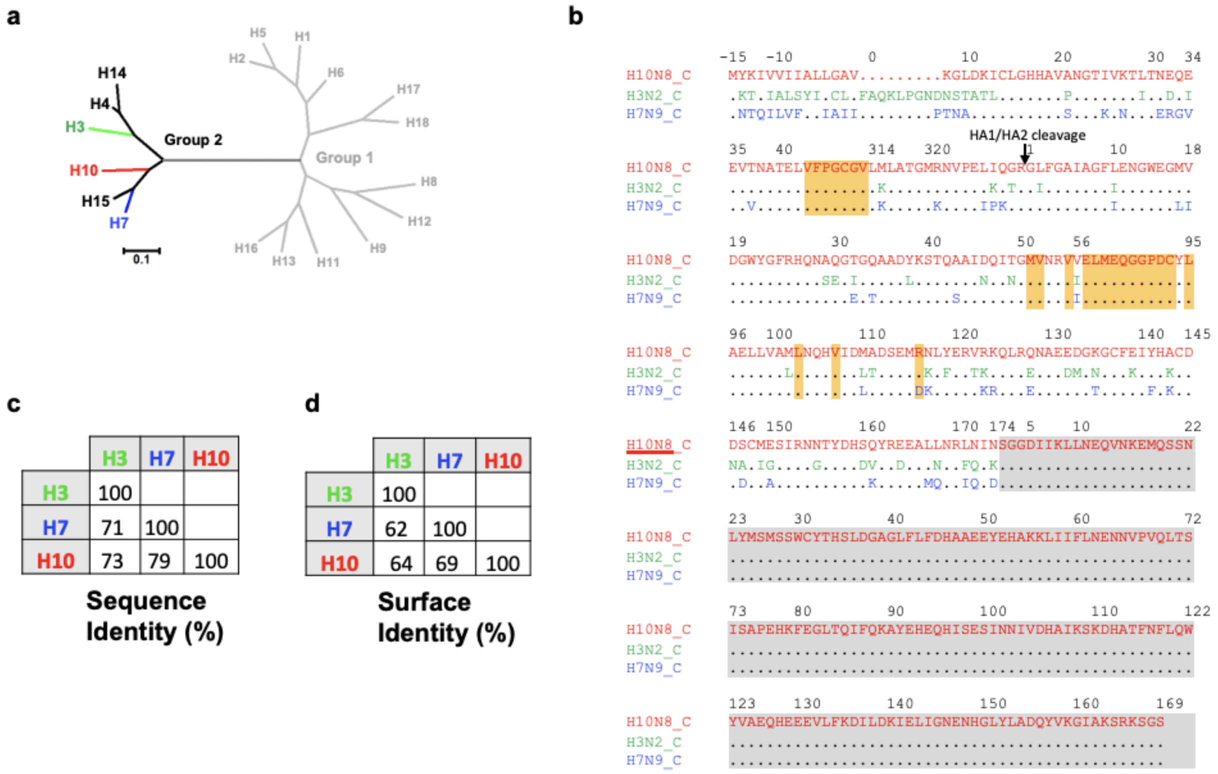


**Supplementary Fig. 1 |** Sequence comparison of group 2 HA stem nanoparticle sequences. **a,** Maximum likelihood phylogenetic tree for protein sequences from representative HA stem of 18 influenza A virus HA subtypes. Group 1 and group 2 HA subtypes are shown in gray and black, respectively. The seasonal (H3) and potential pandemic strains (H7 and H10) are colored in green, blue, and red, respectively. **b,** Sequence alignment of H10ssF, H3ssF, and H7ssF. Sequences were aligned with MUSCLE[75] and each HA subtype color-coded as in **a**. Deletions are indicated in dashes and residues identical to H10ssF are shown in dots. Residues incorporated specifically for stabilization of the stem are shaded in orange while ferritin and linker residues are shaded in gray. HA residues are numbered according to H3N2 HA numbering[76] and ferritin residues are numbered according to PDB entry 3EGM. Amino acid identities of sequence (**c**) and surface (**d**) between H10, H3 and H7 stem.

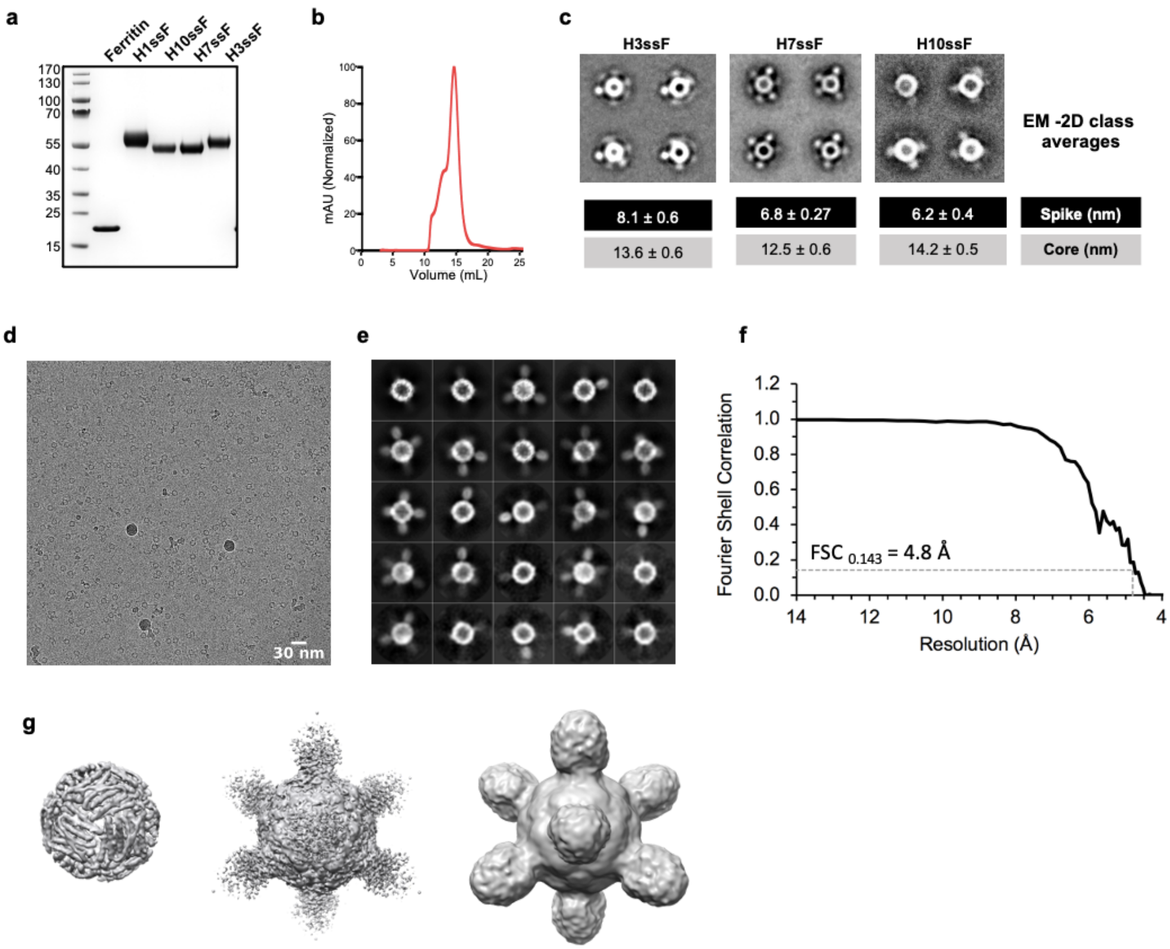


**Supplementary Fig. 2 |** Structural characterization of Group 2 HAssF.

**a,** Expression of HAssF analyzed by SDS-PAGE. Samples were reduced and heat denatured. Molecular weights were denoted on the left margin (kDa). **b,** Size exclusion chromatograms of purified H10ssF. GNA lectin-purified H10ssF was loaded on a Superose 6 10/300 GL column. **c,** Negative stain EM analysis of group 2 HAssF. Representative 2D class averages of each construct are shown. **d,** Raw cryo-EM micrographs of H10ssF. **e,** Representative 2D class averages of H10ssF used in 3D reconstruction. Only a subset of the 8 HA spikes per particle are visible in each class average, consistent with spike flexibility. **f,** Fourier shell correlation (FSC) curve of the H10ssF. A 0.143 cutoff was used for a "gold-standard" refinement indicating a resolution of 4.8 Å for the 3D reconstruction. **g,** Reconstructed 3D model of H10ssF. At a high (left) and a low (middle) contour level in the electron density map were shown. Low-pass filtering the map to 8 Å (right) was shown.

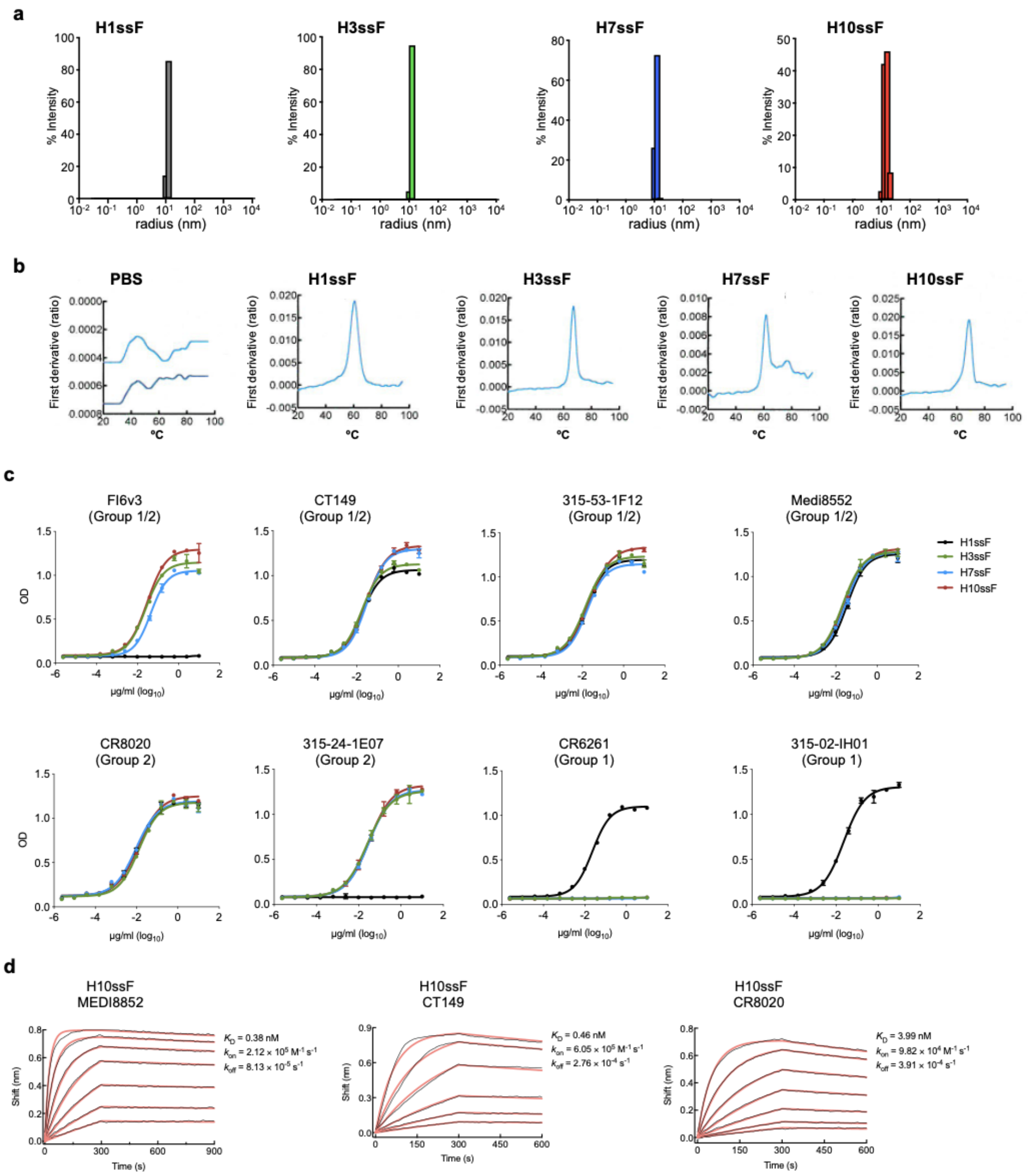


**Supplementary Fig. 3 |** Biophysical and antigenic characterization of group 2 HAssF.

**a,** Dynamic light scattering (DLS) profile of HAssF. Analysis results in a homogeneous species with a radius of ~12nm. **b,** Nano differential scanning fluorometry (nDSF) profile of HAssF. $T_m$ of various HAssF was determined. **c,** Antigenicity of HAssF. ELISA binding curve of stem-specific mAbs to HAssF. FI6v3, CT149, 315-53-1F12 and MEDI8552 were used as cross-group binding mAbs. CR8020 and 315-24-1E07 are specific to group 2 HA stem. CR6261 and 315-02-1H01 are specific to group 1 HA stem. **d,** Binding kinetics of H10ssF to stem-specific Fabs were determined by biolayer interferometry (BLI). Measured sensorgram and calculated curve fit are represented in black and red, respectively. Experiments were performed independently 3 times with similar results.

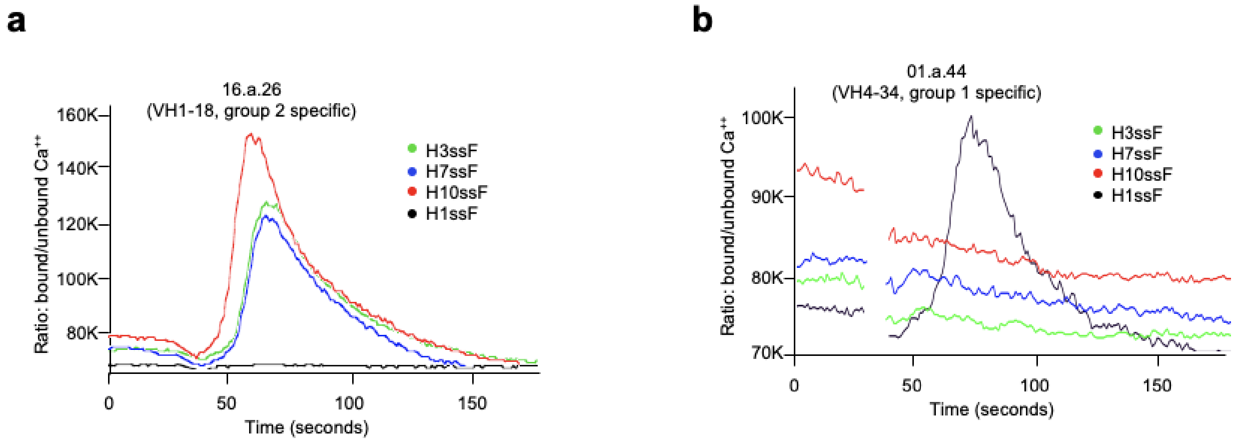


**Supplementary Fig. 4 |** Activation of Ramos B cells expressing HA stem-specific BCRs by HAssF. HAssF were assessed for their ability to activate recombinant Ramos B cells expressing the UCA of stem-specific human B cell receptors. **a,** $Ca^{2+}$ flux profile of group 2 stem-specific BCR of 16.a.26. H3ssF, H7ssF and H10ssF but not group 1 H1ssF activated this BCR. **b,** $Ca^{2+}$ flux profile of group 2 stem-specific BCR of 01.a.44. H1ssF but not group 2 H3ssF, H7ssF and H10ssF activated this BCR.

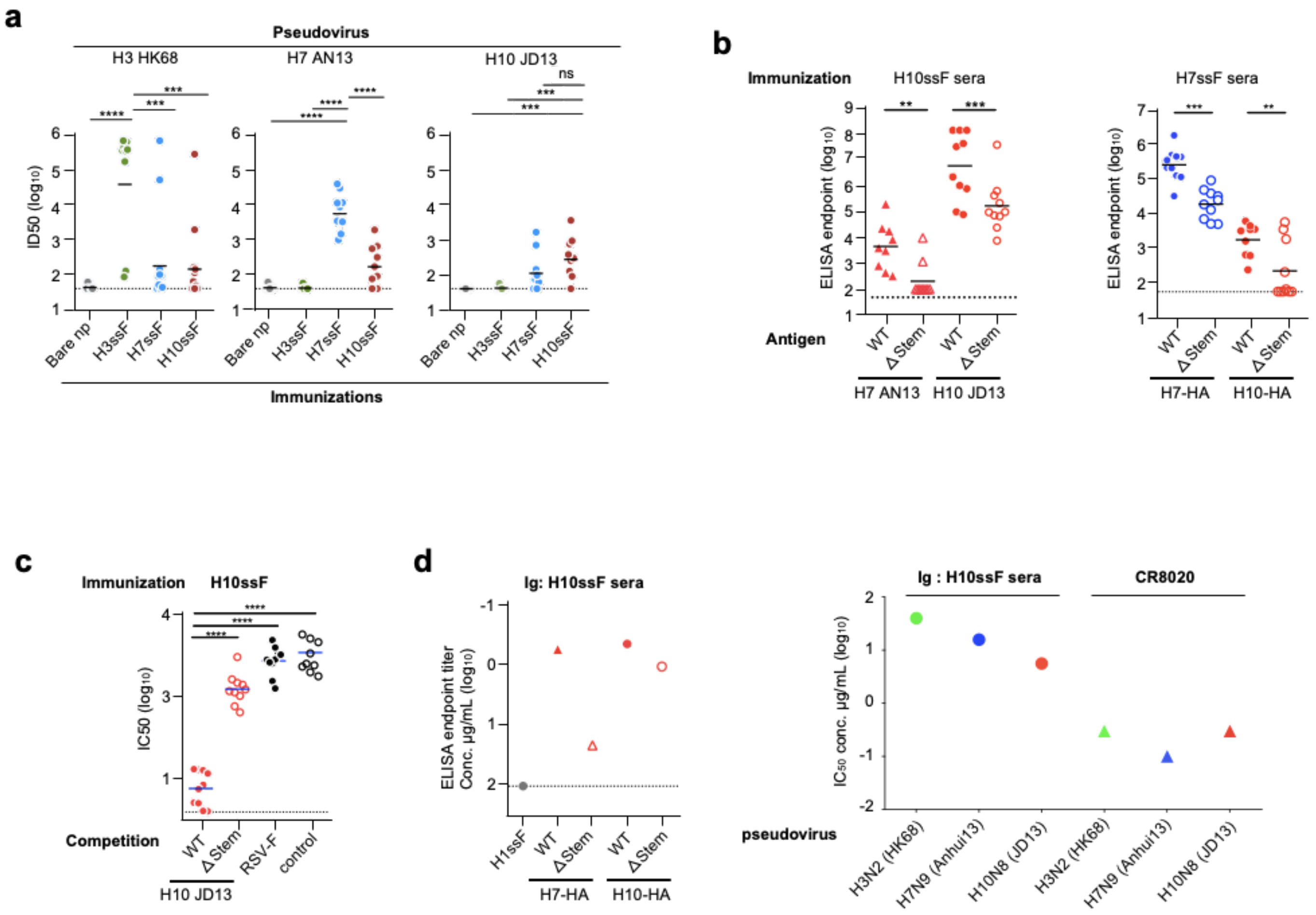


**Supplementary Fig. 5 |** Specificity of group 2 HAssF-induced antibody responses in mice.
**a,** Serum pseudovirus neutralizing antibody titers in group 2 HAssF-immunized mice. Sera of H3ssF-, H7ssF-, H10ssF-, or bare ferritin nanoparticle (bare np)-immunized BALB/cJ mice (n = 10) were tested for neutralization against A/Hong Kong/1/1968 (H3N2), A/Anhui/1/2013 (H7N9), and A/Jiangxi-Donghu/346/13 (H10N8) pseudoviruses. Serum $IC_{50}$ titers were calculated from neutralization curves and plotted. **b,** Serum antibody binding profile to HA and Δstem HA. Immune sera obtained from mice immunized with H10ssF or H7ssF (n = 10) were tested for binding to H7 AN13 HA, H10 JD13 HA, and their Δstem HA with an extra N-glycosylation at the residue 45 in HA2 which knocks out binding of most stem-directed antibodies[26]. **c,** Neutralization competition profile of the H10ssF-immune sera. Excess amounts of H10 JD13 HA and its Δstem HA were used as competitors in the pseudovirus neutralization assays. Respiratory syncytial virus fusion protein (RSV-F) was used as an irrelevant competitor for the assays. **d,** Binding and neutralization profiles of hyper immune immunoglobulins (Igs) isolated from H10ssF-immunized mice. Hyper immune Igs were purified from pooled sera. Binding to H7 AN13 HA, H10 JD13 HA, and their Δstem HA was assessed by ELISA (left). Neutralizing activity of the hyper immune Igs against H3N2, H7N9, and H10N8 pseudoviruses. Horizontal lines indicate geometric means for each group (a-c) and dotted lines indicate lower level of detection (a-d).

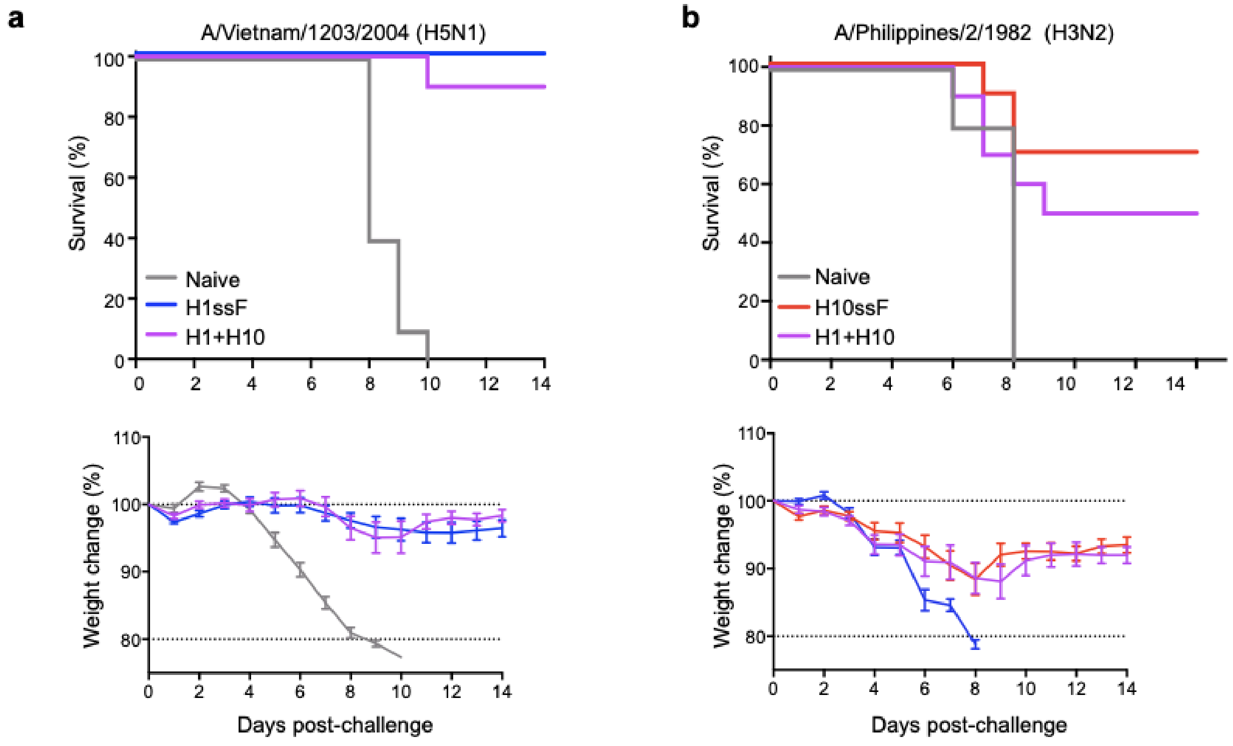


**Supplementary Fig. 6 |** Protection against heterosubtypic lethal virus challenges conferred by H1+10ssF co-immunization in mice.

BALB/cJ mice (n = 10) were co-immunized with 3 doses of 2 µg each of H1ssF and H10ssF at weeks 0, 4, and 8. Ten weeks after the last immunization, mice were experimentally infected with a lethal dose of H5N1 A/Vietnam/1203/2004 (**a**) or H3N2 A/Philippines/2/1982 (**b**). H1ssF- or H10ssF-immunized group served as a control for H5N1 or H3N2 challenges, respectively. Weight loss was recorded for 14 days post infection, animals were euthanized when losing >20% of initial body weight.

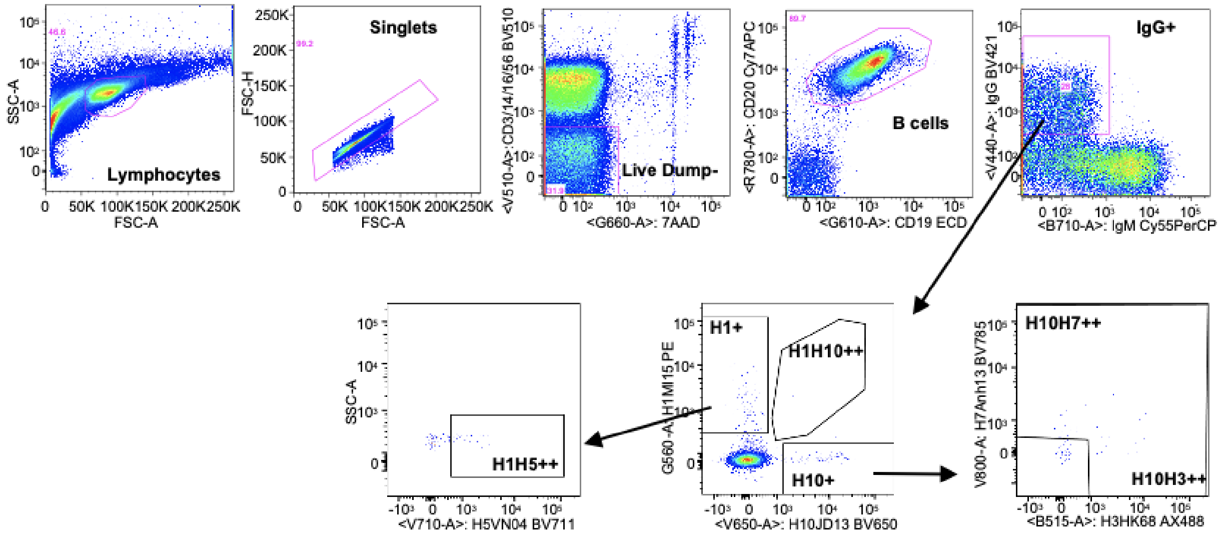


**Supplementary Fig. 7 |** Flow cytometry gating scheme for HA-specific B cells.

To enrich B cells, CD14⁺, CD16⁺, CD56⁺ and CD3⁺ cells were gated out. B cells were further gated for CD19⁺, CD20⁺, IgM⁻, and IgG⁺ prior to applying antigen specific gates. Naïve NHP PBMCs were used to set the threshold for HA probes.

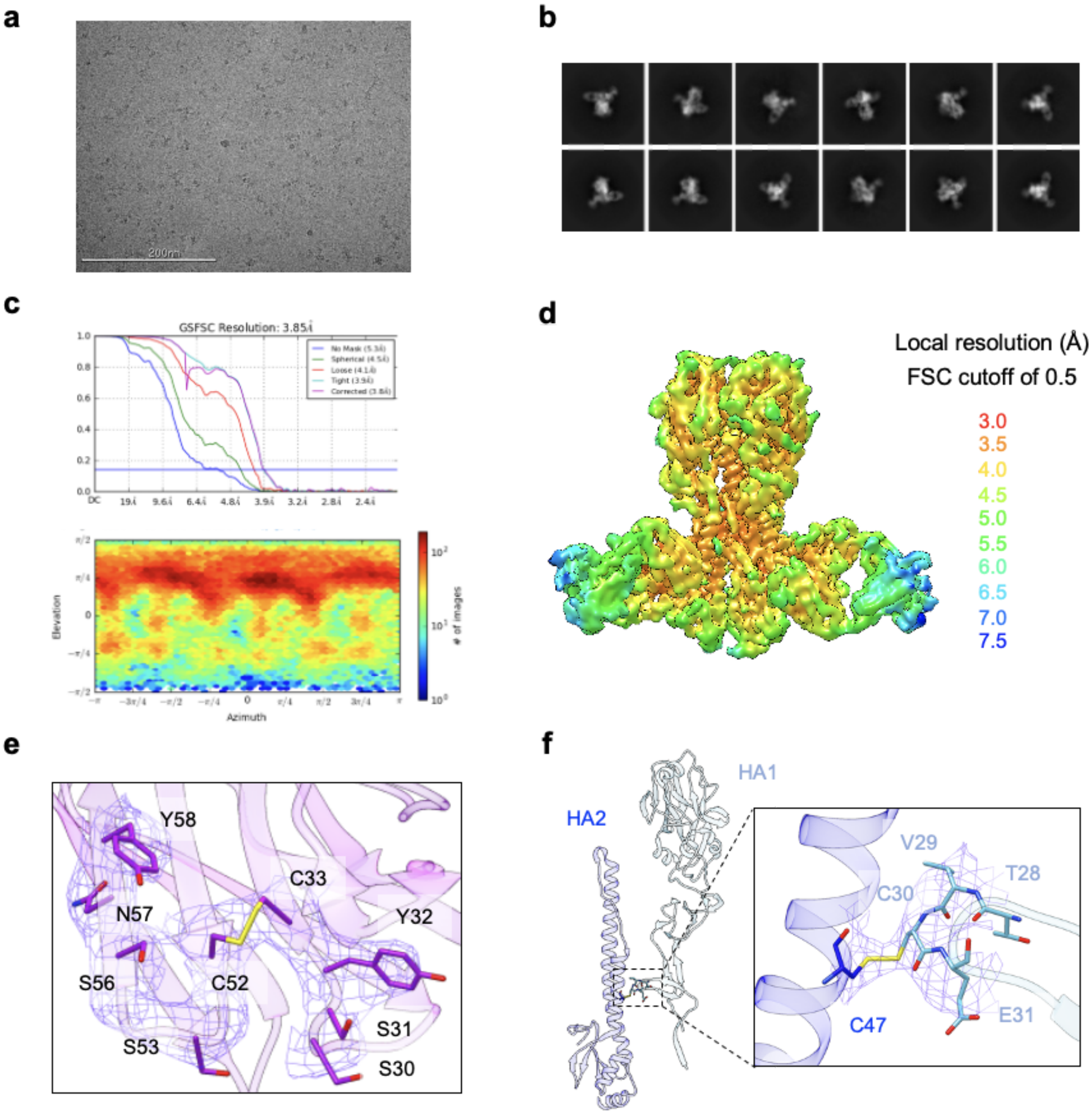


**Supplementary Fig. 8 |** Cryo-EM of 789-203-3C12 Fab in complex with H1 SI06 HA trimer.

**a,** Representative micrograph. **b,** Representative 2D class averages. **c,** The gold-standard FSC resulted in a resolution of 3.85 Å using non-uniform refinement (top); the orientations of all particles used in the final refinement are shown as a heatmap (bottom). **d,** The local resolution of the final map, generated through cryoSPARC using an FSC cutoff of 0.5. **e,** Representative density for the disulfide bond between Cys33 and Cys52 and the surrounding residues in the heavy chain of 789-203-3C12 Fab. Carbon atoms are colored in purple, sulfur in yellow, oxygen in red, nitrogen in blue. **f,** Representative density for the disulfide bond between $Cys47_{HA2}$ and $Cys30_{HA1}$. The HA1 (light blue) and HA2 (blue) protein chains linked by the disulfide bond belong to two different protomers of the trimer. HA1 carbon atoms are colored in light blue, HA2 carbon atoms are colored in blue, sulfur in yellow, oxygen in red, nitrogen in blue.

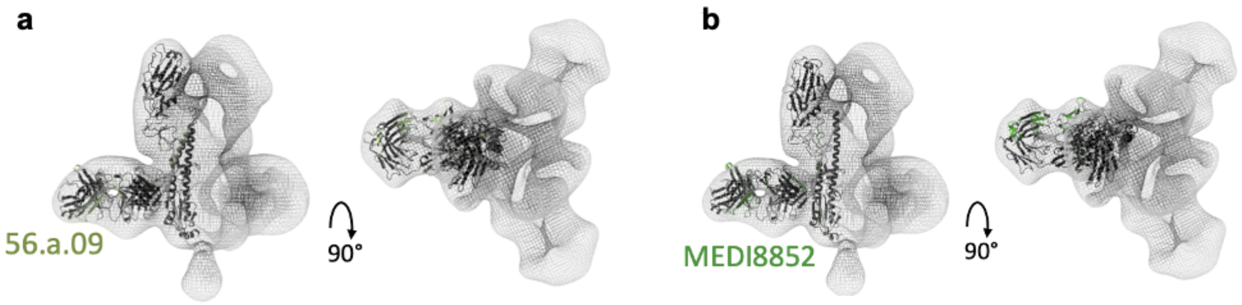


**Supplementary Fig. 9 |** Negative stain EM reconstruction of 789-203-3C12 Fab in complex with H10 JD13 HA.

The density map (mesh) was used to fit crystal structure of 56.a.09–HA (H3, PDB: 5K9K) (**a**) and MEDI8852–HA (H7, PDB: 5JW3) (**b**). Side view with 2 visible Fabs (left) and top view with 3 Fabs (right) are shown.

### Supplementary Table 1. Cryo-EM Data Collection and Refinement Statistics

| | 789-203-3C12 Fab in complex with H1 SI06 HA trimer |
|---|---|
| **EMDB ID** | EMD-22302 |
| **PDB ID** | 6XSK |
| Data Collection | |
| Microscope | FEI Titan Krios |
| Voltage (kV) | 300 |
| Electron dose ($e^-/Å^2$) | 41.92 |
| Detector | Gatan K3 BioQuantum |
| Pixel Size (Å) | 1.07 |
| Defocus Range (μm) | -0.8/-2.5 |
| Magnification | 81000 |
| Reconstruction | |
| Software | cryoSPARC v2.14 |
| Particles | 116,041 |
| Symmetry | C1 |
| Box size (pix) | 360 |
| Resolution (Å) ($FSC_{0.143}$) | 3.85 |
| Refinement | |
| Software | Phenix 1.18 |
| Protein residues | 2163 |
| Chimera CC | 0.875 |
| EMRinger Score | 1.60 |
| R.m.s. deviations | |
| Bond lengths (Å) | 0.004 |
| Bond angles (°) | 0.839 |
| Validation | |
| Molprobity score | 1.35 |
| Clash score | 4.61 |
| Favored rotamers (%) | 100 |
| Ramachandran | |
| Favored regions (%) | 97.4 |
| Allowed regions (%) | 2.6 |
| Disallowed regions (%) | 0 |

### Supplementary Table 2. Interaction Between 789-203-3C12 and H1 SI06 HA

| **Epitope and Paratope Areas** | | |
|---|---|---|
| | HA | Light chain |
| Interface (Å²) | 270.08 | 269.15 |
| | HA | Heavy chain |
| Interface (Å²) | 684.05 | 797.17 |

**Supplementary Table 3. Hydrogen Bond and Salt Bridge Interaction Between 789-203-3C12 and SI06 HA**

| **Hydrogen bonds** | | | |
|---|---|---|---|
| | SI06 HA2 | Bond distance [Å] | 789-203-3C12 Fab |
| 1 | TYR 34 [ N ] | 2.7 | GLY 55 [O ] |
| 2 | GLN 42 [ NE2] | 2.9 | SER 91 [ O ] |
| 3 | GLN 42 [ OE1] | 3 | THR 92 [ OG1] |
| 4 | ASN 46 [ OD1] | 2.9 | ARG 50 [ NH2] |
| 5 | THR 49 [ OG1] | 3.3 | ARG 50 [ NH1] |

| **Salt bridges** | | | |
|---|---|---|---|
| | SI06 HA2 | Distance [Å] | 789-203-3C12 Fab |
| 1 | ASP 19 [ OD2] | 3.6 | ARG 100F[ NH1] |
| 2 | ASP 19 [ OD1] | 3.7 | ARG 100F[ NH2] |
| 3 | ASP 19 [ OD2] | 2.9 | ARG 100F[ NH2] |
| 4 | ASP 146 [ OD2] | 3.2 | LYS 64 [ NZ ] |

### Supplementary Table 4. SI06 HA Residues That Interact with 789-203-3C12 Heavy Chain

<table>
<tr><th>SI06</th><th>HA1</th><th>HSDC</th><th>ASA</th><th>BSA</th><th>ΔiG</th></tr>
<tr><td>8</td><td>C:HIS 18</td><td></td><td>114.7</td><td>22.66 ||</td><td>0.09</td></tr>
<tr><td>[ CB ]</td><td></td><td></td><td></td><td></td><td></td></tr>
<tr><td>[ CG ]</td><td></td><td></td><td></td><td></td><td></td></tr>
<tr><td>[ ND1]</td><td></td><td></td><td></td><td></td><td></td></tr>
<tr><td>[ CD2]</td><td></td><td></td><td></td><td></td><td></td></tr>
<tr><td>[ CE1]</td><td></td><td></td><td></td><td></td><td></td></tr>
<tr><td>[ NE2]</td><td></td><td></td><td></td><td></td><td></td></tr>
<tr><td>10</td><td>C:ASN 20</td><td></td><td>74.34</td><td>19.24 |||</td><td>-0.25</td></tr>
<tr><td>[ OD1]</td><td></td><td></td><td></td><td></td><td></td></tr>
<tr><td>[ ND2]</td><td></td><td></td><td></td><td></td><td></td></tr>
<tr><td>27</td><td>C:THR 37</td><td></td><td>14.54</td><td>7.04 |||||</td><td>0</td></tr>
<tr><td>[ C ]</td><td></td><td></td><td></td><td></td><td></td></tr>
<tr><td>[ O ]</td><td></td><td></td><td></td><td></td><td></td></tr>
<tr><td>[ CB ]</td><td></td><td></td><td></td><td></td><td></td></tr>
<tr><td>[ OG1]</td><td></td><td></td><td></td><td></td><td></td></tr>
<tr><td>28</td><td>C:HIS 38</td><td></td><td>107.3</td><td>50.10 |||||</td><td>0.53</td></tr>
<tr><td>[ CB ]</td><td></td><td></td><td></td><td></td><td></td></tr>
<tr><td>[ CG ]</td><td></td><td></td><td></td><td></td><td></td></tr>
<tr><td>[ ND1]</td><td></td><td></td><td></td><td></td><td></td></tr>
<tr><td>[ CD2]</td><td></td><td></td><td></td><td></td><td></td></tr>
<tr><td>[ CE1]</td><td></td><td></td><td></td><td></td><td></td></tr>
<tr><td>[ NE2]</td><td></td><td></td><td></td><td></td><td></td></tr>
<tr><td>30</td><td>C:VAL 40</td><td></td><td>69.97</td><td>14.05 |||</td><td>0.22</td></tr>
<tr><td>[ CG2]</td><td></td><td></td><td></td><td></td><td></td></tr>
<tr><td>315</td><td>C:THR 318</td><td></td><td>80.62</td><td>29.49 ||||</td><td>0.32</td></tr>
<tr><td>[ C ]</td><td></td><td></td><td></td><td></td><td></td></tr>
<tr><td>[ O ]</td><td></td><td></td><td></td><td></td><td></td></tr>
<tr><td>[ CB ]</td><td></td><td></td><td></td><td></td><td></td></tr>
<tr><td>[ OG1]</td><td></td><td></td><td></td><td></td><td></td></tr>
<tr><td>[ CG2]</td><td></td><td></td><td></td><td></td><td></td></tr>
<tr><td>316</td><td>C:GLY 319</td><td></td><td>24.37</td><td>2.41 |</td><td>0.04</td></tr>
<tr><td>[ CA ]</td><td></td><td></td><td></td><td></td><td></td></tr>
<tr><td>324</td><td>C:NAG 515</td><td></td><td>132.3</td><td>8.50 |</td><td>0</td></tr>
<tr><td>[ O3 ]</td><td></td><td></td><td></td><td></td><td></td></tr>
<tr><td>[ O7 ]</td><td></td><td></td><td></td><td></td><td></td></tr>
</table>

H: Hydrogen bond, S: Salt bridge,
ASA: Accessible Surface Area, Å²,
BSA: Buried Surface Area, Å²,
ΔiG: Solvation energy effect.

<table>
<tr><th>SI06</th><th>HA2</th><th>HSDC</th><th>ASA</th><th>BSA</th><th>ΔiG</th></tr>
<tr><td>10</td><td>B:GLY 16</td><td></td><td>52.02</td><td>21.31 |||||</td><td>-0.14</td></tr>
<tr><td>[ CA ]</td><td></td><td></td><td></td><td></td><td></td></tr>
<tr><td>[ C ]</td><td></td><td></td><td></td><td></td><td></td></tr>
<tr><td>[ O ]</td><td></td><td>H</td><td></td><td></td><td></td></tr>
<tr><td>11</td><td>B:MET 17</td><td></td><td>32.42</td><td>11.21 ||||</td><td>0.18</td></tr>
<tr><td>[ CA ]</td><td></td><td></td><td></td><td></td><td></td></tr>
<tr><td>[ CB ]</td><td></td><td></td><td></td><td></td><td></td></tr>
<tr><td>12</td><td>B:VAL 18</td><td></td><td>136.9</td><td>97.40 ||||||||</td><td>0.92</td></tr>
<tr><td>[ N ]</td><td></td><td></td><td></td><td></td><td></td></tr>
<tr><td>[ O ]</td><td></td><td>H</td><td></td><td></td><td></td></tr>
<tr><td>[ CB ]</td><td></td><td></td><td></td><td></td><td></td></tr>
<tr><td>[ CG1]</td><td></td><td></td><td></td><td></td><td></td></tr>
<tr><td>[ CG2]</td><td></td><td></td><td></td><td></td><td></td></tr>
<tr><td>13</td><td>B:ASP 19</td><td></td><td>95.77</td><td>94.05 ||||||||||</td><td>-0.12</td></tr>
<tr><td>[ N ]</td><td></td><td></td><td></td><td></td><td></td></tr>
<tr><td>[ CA ]</td><td></td><td></td><td></td><td></td><td></td></tr>
<tr><td>[ O ]</td><td></td><td></td><td></td><td></td><td></td></tr>
<tr><td>[ CB ]</td><td></td><td></td><td></td><td></td><td></td></tr>
<tr><td>[ CG ]</td><td></td><td></td><td></td><td></td><td></td></tr>
<tr><td>[ OD1]</td><td></td><td>S</td><td></td><td></td><td></td></tr>
<tr><td>[ OD2]</td><td></td><td>S</td><td></td><td></td><td></td></tr>
<tr><td>14</td><td>B:GLY 20</td><td></td><td>9.04</td><td>5.65 |||||||</td><td>0.09</td></tr>
<tr><td>[ CA ]</td><td></td><td></td><td></td><td></td><td></td></tr>
<tr><td>[ C ]</td><td></td><td></td><td></td><td></td><td></td></tr>
<tr><td>15</td><td>B:TRP 21</td><td></td><td>134.9</td><td>49.30 ||||</td><td>0.37</td></tr>
<tr><td>[ N ]</td><td></td><td></td><td></td><td></td><td></td></tr>
<tr><td>[ CG ]</td><td></td><td></td><td></td><td></td><td></td></tr>
<tr><td>[ CD1]</td><td></td><td></td><td></td><td></td><td></td></tr>
<tr><td>[ CD2]</td><td></td><td></td><td></td><td></td><td></td></tr>
<tr><td>[ NE1]</td><td></td><td></td><td></td><td></td><td></td></tr>
<tr><td>[ CE2]</td><td></td><td></td><td></td><td></td><td></td></tr>
<tr><td>[ CZ2]</td><td></td><td></td><td></td><td></td><td></td></tr>
<tr><td>[ CH2]</td><td></td><td></td><td></td><td></td><td></td></tr>
<tr><td>20</td><td>B:HIS 26</td><td></td><td>43.61</td><td>1.72 |</td><td>0.03</td></tr>
<tr><td>[ CE1]</td><td></td><td></td><td></td><td></td><td></td></tr>
<tr><td>26</td><td>B:SER 32</td><td></td><td>70.08</td><td>3.07 |</td><td>-0.04</td></tr>
<tr><td>[ O ]</td><td></td><td></td><td></td><td></td><td></td></tr>
<tr><td>27</td><td>B:GLY 33</td><td></td><td>25.29</td><td>13.91 ||||||</td><td>0.22</td></tr>
<tr><td>[ CA ]</td><td></td><td></td><td></td><td></td><td></td></tr>
<tr><td>28</td><td>B:TYR 34</td><td></td><td>107.6</td><td>50.77 |||||</td><td>0.17</td></tr>
<tr><td>[ N ]</td><td></td><td>H</td><td></td><td></td><td></td></tr>
<tr><td>[ C ]</td><td></td><td></td><td></td><td></td><td></td></tr>
<tr><td>[ O ]</td><td></td><td></td><td></td><td></td><td></td></tr>
<tr><td>[ CB ]</td><td></td><td></td><td></td><td></td><td></td></tr>
<tr><td>[ CD2]</td><td></td><td></td><td></td><td></td><td></td></tr>
<tr><td>[ CE2]</td><td></td><td></td><td></td><td></td><td></td></tr>
<tr><td>29</td><td>B:ALA 35</td><td></td><td>27.53</td><td>17.50 |||||||</td><td>0.28</td></tr>
<tr><td>[ CA ]</td><td></td><td></td><td></td><td></td><td></td></tr>
<tr><td>[ C ]</td><td></td><td></td><td></td><td></td><td></td></tr>
<tr><td>[ CB ]</td><td></td><td></td><td></td><td></td><td></td></tr>
<tr><td>30</td><td>B:ALA 36</td><td></td><td>23.36</td><td>7.73 ||||</td><td>-0.04</td></tr>
<tr><td>[ N ]</td><td></td><td>H</td><td></td><td></td><td></td></tr>
<tr><td>[ O ]</td><td></td><td></td><td></td><td></td><td></td></tr>
<tr><td>[ CB ]</td><td></td><td></td><td></td><td></td><td></td></tr>
<tr><td>32</td><td>B:GLN 38</td><td></td><td>97.46</td><td>40.17 |||||</td><td>-0.33</td></tr>
<tr><td>[ CB ]</td><td></td><td></td><td></td><td></td><td></td></tr>
<tr><td>[ CD ]</td><td></td><td></td><td></td><td></td><td></td></tr>
<tr><td>[ OE1]</td><td></td><td></td><td></td><td></td><td></td></tr>
<tr><td>[ NE2]</td><td></td><td></td><td></td><td></td><td></td></tr>
<tr><td>35</td><td>B:THR 41</td><td></td><td>4.55</td><td>3.38 ||||||||</td><td>0.04</td></tr>
<tr><td>[ OG1]</td><td></td><td></td><td></td><td></td><td></td></tr>
<tr><td>[ CG2]</td><td></td><td></td><td></td><td></td><td></td></tr>
<tr><td>36</td><td>B:GLN 42</td><td></td><td>117.5</td><td>19.76 ||</td><td>-0.12</td></tr>
<tr><td>[ CG ]</td><td></td><td></td><td></td><td></td><td></td></tr>
<tr><td>[ NE2]</td><td></td><td></td><td></td><td></td><td></td></tr>
<tr><td>[ CG ]</td><td></td><td></td><td></td><td></td><td></td></tr>
<tr><td>[ NE2]</td><td></td><td></td><td></td><td></td><td></td></tr>
<tr><td>39</td><td>B:ILE 45</td><td></td><td>80.51</td><td>55.86 |||||||</td><td>0.89</td></tr>
<tr><td>[ O ]</td><td></td><td></td><td></td><td></td><td></td></tr>
<tr><td>[ CG1]</td><td></td><td></td><td></td><td></td><td></td></tr>
<tr><td>[ CG2]</td><td></td><td></td><td></td><td></td><td></td></tr>
<tr><td>[ CD1]</td><td></td><td></td><td></td><td></td><td></td></tr>
<tr><td>42</td><td>B:ILE 48</td><td></td><td>29.96</td><td>9.54 ||||</td><td>0.15</td></tr>
<tr><td>[ CB ]</td><td></td><td></td><td></td><td></td><td></td></tr>
<tr><td>[ CG2]</td><td></td><td></td><td></td><td></td><td></td></tr>
<tr><td>43</td><td>B:THR 49</td><td></td><td>67.72</td><td>18.40 |||</td><td>0.1</td></tr>
<tr><td>[ N ]</td><td></td><td></td><td></td><td></td><td></td></tr>
<tr><td>[ CA ]</td><td></td><td></td><td></td><td></td><td></td></tr>
<tr><td>[ OG1]</td><td></td><td></td><td></td><td></td><td></td></tr>
<tr><td>[ CG2]</td><td></td><td></td><td></td><td></td><td></td></tr>
<tr><td>46</td><td>B:VAL 52</td><td></td><td>72.14</td><td>17.75 |||</td><td>0.28</td></tr>
<tr><td>[ CB ]</td><td></td><td></td><td></td><td></td><td></td></tr>
<tr><td>[ CG1]</td><td></td><td></td><td></td><td></td><td></td></tr>
<tr><td>[ CG2]</td><td></td><td></td><td></td><td></td><td></td></tr>
<tr><td>140</td><td>B:ASP 146</td><td></td><td>74.48</td><td>20.90 |||</td><td>-0.14</td></tr>
<tr><td>[ CB ]</td><td></td><td></td><td></td><td></td><td></td></tr>
<tr><td>[ CG ]</td><td></td><td></td><td></td><td></td><td></td></tr>
<tr><td>[ OD2]</td><td></td><td>S</td><td></td><td></td><td></td></tr>
<tr><td>144</td><td>B:GLU 150</td><td></td><td>80.69</td><td>7.58 |</td><td>0.12</td></tr>
<tr><td>[ CB ]</td><td></td><td></td><td></td><td></td><td></td></tr>
<tr><td>[ CG ]</td><td></td><td></td><td></td><td></td><td></td></tr>
</table>

### Supplementary Table 5. 789-203-3C12 Heavy Chain Residues That Interact with SI06 HA

| | HC | HSDC | ASA | BSA | ΔiG |
|---|---|---|---|---|---|
| 31 | I:SER 31 | | 91.85 | 25.24 \|\|\| | 0.21 |
| [ CA ] | | | | | |
| [ C ] | | | | | |
| [ O ] | | | | | |
| [ CB ] | | | | | |
| [ OG ] | | | | | |
| 33 | I:CYS 33 | | 5.63 | 5.46 \|\|\|\|\|\|\|\|\|\| | 0.22 |
| [ SG ] | | | | | |
| 52 | I:CYS 52 | | 20.2 | 18.82 \|\|\|\|\|\|\|\|\|\| | 0.75 |
| [ C ] | | | | | |
| [ CB ] | | | | | |
| [ SG ] | | | | | |
| 53 | I:GLY 52A | | 26.82 | 14.21 \|\|\|\|\|\| | 0.22 |
| [ N ] | | | | | |
| [ CA ] | | | | | |
| 55 | I:GLY 54 | | 46.18 | 5.38 \|\| | -0.06 |
| [ O ] | | | | | |
| 56 | I:GLY 55 | | 70.04 | 54.07 \|\|\|\|\|\|\|\| | 0.03 |
| [ CA ] | | | | | |
| [ C ] | | | | | |
| [ O ] | | H | | | |
| 57 | I:SER 56 | | 43.92 | 43.03 \|\|\|\|\|\|\|\|\|\| | -0.24 |
| [ CA ] | | | | | |
| [ CB ] | | | | | |
| [ OG ] | | | | | |
| 58 | I:ASN 57 | | 80.17 | 11.92 \|\| | -0.09 |
| [ N ] | | | | | |
| [ O ] | | | | | |
| [ CB ] | | | | | |
| 59 | I:TYR 58 | | 104.87 | 72.92 \|\|\|\|\|\|\| | 0.2 |
| [ CD1] | | | | | |
| [ CE1] | | | | | |
| [ CE2] | | | | | |
| [ CZ ] | | | | | |
| [ OH ] | | H | | | |
| 65 | I:LYS 64 | | 124.8 | 26.92 \|\|\| | -0.14 |
| [ CG ] | | | | | |
| [ CD ] | | | | | |
| [ CE ] | | | | | |
| [ NZ ] | | | | | |
| 103 | I:ILE 99 | | 101.53 | 22.09 \|\|\| | 0.35 |
| [ CG1] | | | | | |
| [ CD1] | | | | | |
| 104 | I:PHE 100 | | 219.68 | 123.10 \|\|\|\|\|\| | 1.62 |

H: Hydrogen bond, S: Salt bridge,
ASA: Accessible Surface Area, Å²,
BSA: Buried Surface Area, Å²,
ΔiG: Solvation energy effect.

### Supplementary Table 6. SI06 HA Residues That Interact with 789-203-3C12 Light Chain

| SI06 | HA2 | HSDC | ASA | BSA | ΔiG |
|---|---|---|---|---|---|
| 32 | D:GLN 38 | | 92.77 | 36.11 \|\|\|\| | 0.38 |
| [ C ] | | | | | |
| [ CB ] | | | | | |
| [ CG ] | | | | | |
| [ CD ] | | | | | |
| [ NE2] | | | | | |
| 33 | D:LYS 39 | | 140.59 | 20.42 \|\| | 0.33 |
| [ CA ] | | | | | |
| [ CG ] | | | | | |
| [ CE ] | | | | | |
| 36 | D:GLN 42 | | 116.45 | 96.04 \|\|\|\|\|\|\|\|\| | -0.68 |
| [ CA ] | | | | | |
| [ O ] | | | | | |
| [ CB ] | | | | | |
| [ CG ] | | | | | |
| [ CD ] | | | | | |
| [ OE1] | | H | | | |
| [ NE2] | | H | | | |
| 39 | D:ILE 45 | | 87.62 | 31.00 \|\|\|\| | 0.5 |
| [ CB ] | | | | | |
| [ CG2] | | | | | |
| [ CD1] | | | | | |
| 40 | D:ASN 46 | | 106.53 | 62.65 \|\|\|\|\|\| | -0.78 |
| [ CA ] | | | | | |
| [ CG ] | | | | | |
| [ OD1] | | H | | | |
| [ ND2] | | | | | |
| 43 | D:THR 49 | | 67.81 | 23.86 \|\|\|\| | 0.2 |
| [ CB ] | | | | | |
| [ OG1] | | H | | | |
| [ CG2] | | | | | |

**H: Hydrogen bond, S: Salt bridge,**
**ASA: Accessible Surface Area, Å²,**
**BSA: Buried Surface Area, Å²,**
**ΔiG: Solvation energy effect.**

### Supplementary Table 7. 789-203-3C12 Light Chain Residues That Interact with SI06HA

| | LC | HSDC | ASA | BSA | ΔiG |
|---|---|---|---|---|---|
| 29 | J:ILE 29 | | 11.39 | 3.01 \|\|\| | 0.05 |
| [ CG2] | | | | | |
| 30 | J:SER 30 | | 78.66 | 15.77 \|\|\| | 0.21 |
| [ N ] | | | | | |
| [ CB ] | | | | | |
| [ OG ] | | | | | |
| 31 | J:SER 31 | | 49.18 | 7.20 \|\| | 0.12 |
| [ CB ] | | | | | |
| 32 | J:TRP 32 | | 131.32 | 78.46 \|\|\|\|\| | 0.8 |
| [ CB ] | | | | | |
| [ CG ] | | | | | |
| [ CD1] | | | | | |
| [ CD2] | | | | | |
| [ NE1] | | | | | |
| [ CE2] | | | | | |
| [ CE3] | | | | | |
| [ CZ2] | | | | | |
| [ CZ3] | | | | | |
| [ CH2] | | | | | |
| 50 | J:ARG 50 | | 131.7 | 78.60 \|\|\|\|\| | -1.53 |
| [ NE ] | | | | | |
| [ CZ ] | | | | | |
| [ NH1] | | H | | | |
| [ NH2] | | H | | | |
| 91 | J:SER 91 | | 52.54 | 6.26 \|\| | -0.07 |
| [ O ] | | H | | | |
| 92 | J:THR 92 | | 79.88 | 52.34 \|\|\|\|\|\| | -0.33 |
| [ CA ] | | | | | |
| [ C ] | | | | | |
| [ O ] | | | | | |
| [ CB ] | | | | | |
| [ OG1] | | H | | | |
| 93 | J:SER 93 | | 52.07 | 12.28 \|\|\| | 0.13 |
| [ CA ] | | | | | |
| [ CB ] | | | | | |
| [ OG ] | | | | | |
| 96 | J:PHE 96 | | 99.24 | 15.23 \|\| | 0.24 |
| [ CZ ] | | | | | |

**H: Hydrogen bond, S: Salt bridge,**
**ASA: Accessible Surface Area, Å²,**
**BSA: Buried Surface Area, Å²,**
**ΔiG: Solvation energy effect.**
